# Interactions of the LINE-1 encoded ORF1p with proteins and chromatin converge on a role in neuronal physiology

**DOI:** 10.1101/2025.09.05.674414

**Authors:** Sandra Sinnassamy, Olivia Massiani Beaudoin, Berangère Lombard, Damarys Loew, Tom Bonnifet, Magali Fradet, Héloïse Monnet, Thomas Caille, Nicolas Servant, Rajiv L. Joshi, Julia Fuchs

## Abstract

Retrotransposons are emerging as novel regulators of embryonic and brain development. We recently demonstrated that the LINE-1–encoded protein ORF1p is abundantly expressed in adult mouse and human neurons, although its function remains unclear. Here, we characterize the ORF1p interactome in differentiated mouse and human neurons using mass spectrometry and identify novel partners implicated in gene regulation and neuron-specific processes. ORF1p localizes not only to neuronal nuclei, where it associates with chromatin under steady-state conditions, but also to neurites, supporting a role in neuronal physiology. To further explore its nuclear functions, we sorted human post-mortem neurons with high or low nuclear ORF1p levels and performed ORF1p knockdown in cultured human neurons, followed by chromatin accessibility assays. Both approaches revealed consistent patterns of differential chromatin accessibility dependent on ORF1p. Loss of ORF1p also led to the downregulation of long, neuron-specific genes and altered neurite morphology. Together, these findings point to a physiological role of ORF1p in post-mitotic neurons, mediated through converging interactions with proteins and chromatin.

## Introduction

Transposable elements (TEs), once considered junk DNA, are gaining interest as physiological modulators of genome architecture^1^ and early embryonic ^2^ as well as neurodevelopment^3^, but also pathological drivers of aging-related^4^ and neurogenerative processes. Half of the human genome is composed of TEs and 17% of the Long Interspersed Nuclear Element-1 (LINE-1 or L1)^5^ TE family. Between 100 and 150 LINE-1^6,7^ are considered full-length L1 elements (flLINE-1). They are the only autonomous mobile elements in the human genome and are characterized by a bidirectional RNA polymerase II promoter in the 5’UTR and intact open reading frames encoding the canonical L1 proteins, ORF1p and ORF2p, essential for LINE-1 mobilization. ORF1p and ORF2p, translated from a bicistronic mRNA, associate with *cis-preference* to the LINE-1 mRNA to form a ribonucleoprotein (RNP). Once the LINE-1 RNP gets access to the nucleus, a new copy is pasted into the genome through target-primed reverse transcription (TPRT). ORF1p is an RNA binding protein and a nucleic acid chaperon and ORF2p has endonuclease and reverse transcriptase activities. When abnormally expressed, LINE-1 may represent a threat to genome and cellular integrity by generating DNA double strand breaks and triggering inflammatory pathways^8^. Consequently, LINE-1 expression is rigorously controlled through multiple mechanisms at the transcriptional and post-transcriptional levels^9^. Nevertheless, LINE-1 has been shown to exert physiological functions especially during early embryonic^10,11^and neuronal development^3,12–14^. Intriguingly, recent and optimized next generation sequencing data have highlighted the abundant expression of LINE-1 in adult human brain^15,16^, revealing significant enrichment of LINE-1 expression, including evolutionary young L1HS and L1PA2 elements, in adult human neurons with no or limited expression of LINE-1 in non-neuronal cells at steady-state^12,15,16^. Indeed, on the protein level, LINE-1 encoded ORF1p expression was observed predominantly in neurons throughout the adult mouse brain^17^ and in several regions of the human brain^17,18^ as well as in human dopaminergic neurons^17–19^. The presence of ORF1p in adult neurons is a surprising observation given the deleterious consequences to cellular integrity attributed to LINE-1 expression and led us to test the hypothesis of a physiological function of this protein in the adult brain. To this end, we used a combination of mass spectrometry, chromatin accessibility assays and transcriptomics. We found an important protein interaction network of endogenous ORF1p in adult neurons encompassing proteins related to RNA metabolism, as expected for an RNA binding protein, but also proteins related to nuclear functions including chromatin regulation and gene expression as well as proteins with neuron-specific functions. This finding and the presence of ORF1p in chromatin fractions led us to analyse chromatin accessibility in nuclei with high or low ORF1p content using sorted *post-mortem* neurons or ORF1p loss-of-function (ORF1p-LOF) in differentiated human neurons in which we also assessed transcriptomic changes. Characteristics of differentially accessible regions were significantly different between nuclei with high or low ORF1p content and these differences were highly similar across two different experimental systems involving gene loci related to genes important for neuronal functions. ORF1p-LOF led to the downregulation of particularly long, protein-coding genes enriched in neuronal functions. Functionally, we observed morphological changes in neurites of adult mouse neurons upon ORF1p-LOF. Together, these data suggest a role of ORF1p in the maintenance and integrity of neuronal function in adult neurons via converging actions on chromatin organization, gene transcription and protein interactions.

## Results

### Identification of protein partners of endogenous ORF1p in mouse and human neurons

We and others have previously observed a widespread expression of ORF1p in mouse and human neurons at steady-state^17–20^. This somewhat surprising presence of ORF1p in neurons evokes the question of a possible physiological role of ORF1p specific to these cells. In a first attempt to address this question, we identified, using a quantitative mass spectrometry approach, protein partners of endogenous ORF1p in mouse (differentiated MN9D) and human (differentiated LUHMES^19,21^) dopaminergic neurons (Fig. 1A) using well characterized and specific antibodies to ORF1p^17^. Just as differentiated human dopaminergic LUHMES neurons^19,21^, mouse dopaminergic MND9 precursor cells can be differentiated into mature, post-mitotic neurons (Suppl. Fig. 1A) which express ORF1p (Suppl. Fig. 1B) along with the neuronal marker β-III tubulin (TUJ1, Suppl. Fig. 1C) and the dopaminergic marker tyrosine hydroxylase (TH, Suppl. Fig. 1C). As expected, the pattern of ORF1p staining was mainly cytoplasmic, but we also observed, as in human dopaminergic LUHMES^19^, a readily detectable nuclear staining (Suppl. Fig. 1B). It is important to note that human and mouse ORF1p differ significantly in their amino acid composition (31% identity, NCBI Needleman-Wunsch Global Align), but share the same protein domains^22^ which is reflected in very similar Alpha Fold predicted 3D protein structures for mouse^23^ and human^24^ ORF1p. Following immunoprecipitation (IP) of endogenous ORF1p or isotype control IgG in human (LUHMES) and mouse (MN9D) dopaminergic neurons (n=5 IPs per condition), we performed liquid chromatography coupled to mass spectrometry (LC-MS/MS) followed by a quantitative analysis (Fig. 1A). ORF1p was strongly enriched in the immunoprecipitated material, with a high IP-ORF1p/IP-IgG ratio (mouse ORF1p: 3.76 log2(ratio), p= 4.16E-05, Fig. 1B; human ORF1p: 5.35 log2(ratio), p= 8.73E-07, Fig. 1C). 348 proteins were detected as ORF1p partners in MN9D (Fig. 1B) and 267 proteins were identified as ORF1p partner proteins in LUHMES neurons (Fig. 1C). As some proteins detected in the ORF1p IP were completely absent from the control IP-IgG and, as a consequence, no ratio could be calculated, these putative ORF1p protein partners are represented separately as peptides/100aa (Fig. 1B, C, right panels). Next, we confronted ORF1p protein partners identified in neurons with those found in previous studies^25–34^ including our previous study of ORF1p interactors in the mouse brain^17^. Nine proteins were identified in all studies (Suppl. Fig. 1D), namely RPL19, IGF2BP3, LARP1, ATXN2, RPL24, YBX1, DDX6, PABPC1 and RPL8, representing thus high-confidence, highly conserved and non-cell-type specific ORF1p protein partners most of which were RNA binding proteins (RBPs), just as ORF1p itself (Suppl. Fig. 1H, I). 49 proteins were shared between ORF1p partners identified in various human cell types and ORF1p partners identified in human LUHMES neurons (Suppl. Fig. 1E), among them: *HNRNPU, NCL* and *IGF2BP2* and of those 14 proteins were overlapping in addition with ORF1p protein partners identifies in mouse MN9D neurons, including RPL19, IGF2BP3, RALY, LARP, ATXN2, RPL24, YBX1, CLASP1, DDX6, UBAP2L, DDX17, ELAVL1, PABPC1 and RPL8 (Suppl Table. 1). Similarly, 70 proteins among which GPHN, ELAVL2, ARID1B, FXR1, POLR2A, MAP1B, EEA1, ARID1A and HELZ were in common between previously identified mouse ORF1p partners and those identified in MN9D neurons (Suppl. Fig. 1F). Focusing on overlapping ORF1p protein partners in mouse and human neurons (Suppl. Fig. 1G), the 41 proteins that were shared mostly identified as “RNA processing” and/or “RNA binding” in a STRING analysis visualized by Cytoscape (Suppl. Fig. 1G). To identify conserved categories of ORF1p protein partners between mouse and human neurons, we inquired for shared significant GO terms. As expected, among the top shared categories, many GO terms were related to RNA metabolism (Fig. 1D, cat.III = category 3 interactors). However, two unexpected categories emerged: first, in both mouse and human neurons, ORF1p protein partners were enriched in GO terms related to nuclear presence and function (cat.I = category I interactors), namely “gene expression” (107 proteins in LUHMES (fold enrichment (FE): 3,25; FDR:1,61E-26), 76 in MN9D (FE: 1,76; FDR: 0,0001), 23 in common) and “nucleus” (186 proteins in LUHMES (FE: 1,37; FDR: 3,07E-05), 162 in MN9D (FE: 1,82; FDR: 3,47E-22), 23 in common). In this category I of ORF1p protein partners, 31 unique proteins were identified, among which 17 ribosomal proteins with non-canonical functions, in addition to DDX6, FXR1, GSN, IGF2BP3, KHDRBS1, LARP1, MLF2, MYH9, PABPC1, PRKRA, YBX1, HNRNPU, YTHDC1 and RCOR2, the last three of which have chromatin-related functions. In LUHMES cells, ORF1p interacted with proteins related to chromatin organization and epigenetic regulation (H1-2, H1-4, H1-10, MACROH2A1, MACROH2A2, CHD4, CBX8, ASH1L, SUPT16H, DPF2, SMARCA4, MORC2), proteins related to DNA damage and repair (TOP2B, PARP1, TP53BP1, XRCC6 (Ku70) and GADD45GP1), proteins involved in transcriptional regulation (CTNNB1, YBX1, ZC3H11A, THRAP3, SNW1, BCLAF1, PSIP1, TFAP2D), proteins associated nuclear structure and envelope (LMNA, LBR, TMPO, EMD, NUP153, NUP214, RANBP2, NUMA1) and nuclear transport and signalling (PAWR, FXR1 and FXR2) in addition to many more involved in RNA processing, RNA metabolism and ribosome biogenesis (Suppl Table 1). In MN9D neurons, ORF1p interacted with chromatin remodelers (CHD7, ARID1B, ARID1A), transcription factors (ZFHX3, STAT2, ZNF638, ZNF281, TCF4), with RNA polymerase subunits or associated proteins (POLR2M, POLR2A, POLR2B, POLR2C, RPAP2) and nuclear transport proteins (NUP50 and KPNA3) in addition to many RNA metabolism related proteins (Suppl Table 1). In the second category (cat.II interactors), ORF1p protein partners were enriched for GO terms related to neuron morphology like ““synapse” (LUHMES; FE: 1,92; FDR: 001; MN9D; FE: 3,62; FDR:13E-32), “neuron projection” (LUHMES; FE: 1,85; FDR: 0,01; MN9D; FE: 2,46; FDR: 6,35E-10), “dendrite” (LUHMES; FE: 2,15; FDR: 0,0313; MN9D; FE: 3,13; FDR: 1,93E-09), “axo-dendritic transport” (LUHMES; FE: 5,84; FDR: 0,04; MN9D; FE: 5,53; FDR: 0,008), “cytoskeleton protein binding” (LUHMES; FE: 2,73; FDR: 1,12E-05; MN9D; FE: 4,04; FDR: 1,01E-19) and “actin binding” (LUHMES; FE: 3,91; FDR: 7,70E-06; MN9D; FE: 4,91; FDR: 3,10E-12) (Fig. 1D). STRING analysis of physical protein-protein interactions of ORF1p partners in mouse (Fig. 1E) and human (Fig. 1F) neurons revealed the interaction profile of these proteins among each other according to their functional category (colours), confidence in physical interaction (thickness of edges) and known interactors (yellow nodes). The emergence of categories related to nuclear functions, including gene expression, was particularly intriguing in light of observations of a nuclear localization of ORF1p at steady-state in human neurons^19^. Indeed, while ORF1p has been previously observed in the nucleus in different cell lines^28,35–41^ and in human post-mortem neurons^17,18^, the role of nuclear ORF1p has not been previously addressed except with regard to retrotransposition efficiency^42^. We therefore asked the question whether nuclear ORF1p might have a role in mature neurons, potentially independent of the LINE-1 life cycle.

**Figure 1:**
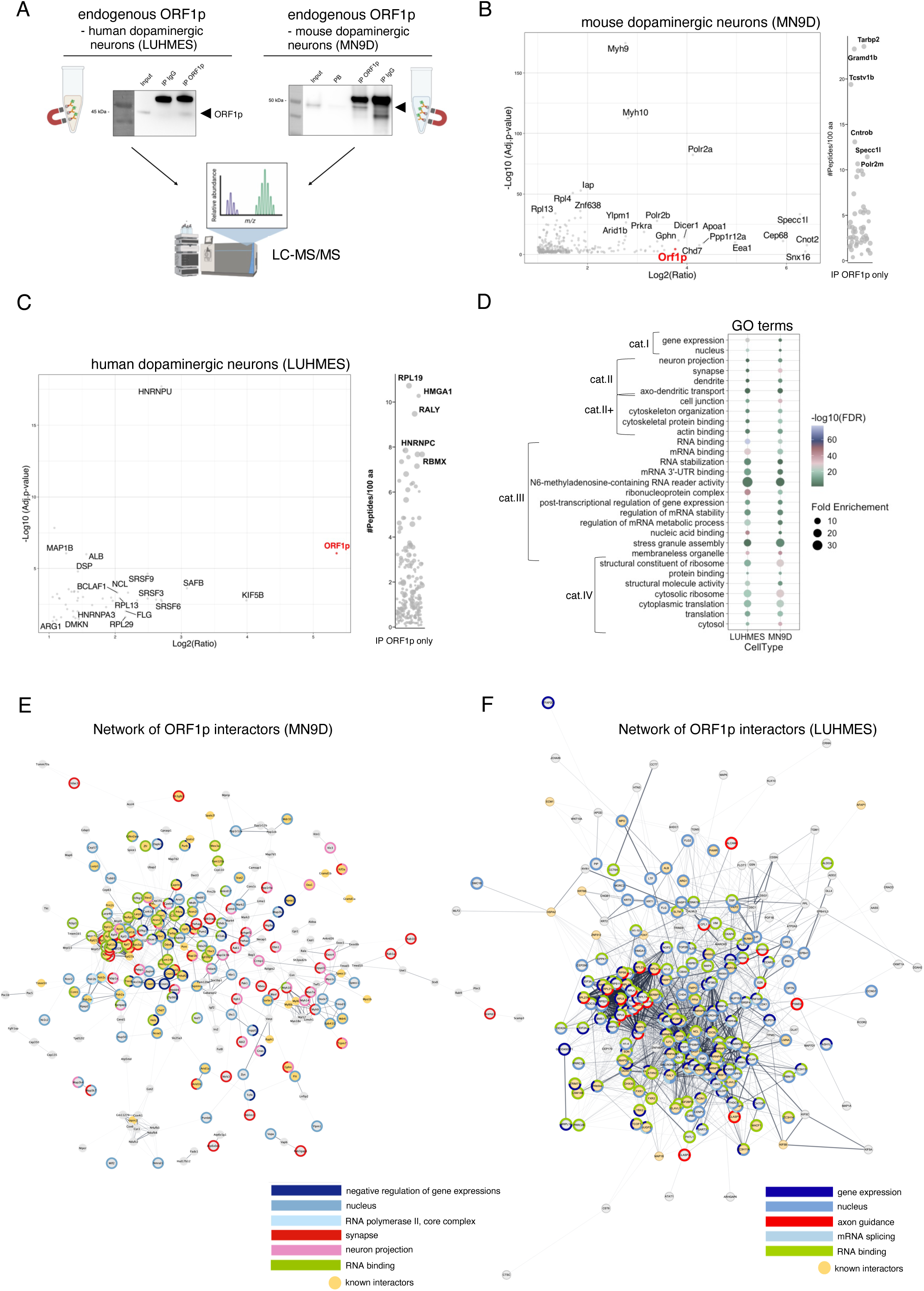
Quantitative mass spectrometry analysis of protein partners of endogenous ORF1p in human and mouse dopaminergic neurons. **A.** Graphical representation of the experimental protocol used to identify protein partners of endogenous ORF1p in mouse and human neurons. Endogenous ORF1p was immunoprecipitated (IP) from human (LUHMES) and mouse (MN9D) differentiated dopaminergic neurons, respectively. Enrichment was verified by Western blot analysis and lysates were analysed by LC-MS/MS followed by the identification and quantification of protein partners. **B.** Specific protein partners of ORF1p found by proteomic mass spectrometry in MN9D neurons, presented according to their fold change (ratio IP-ORF1p/IP-IgG) and adjusted p-value. Proteins identified in the IP ORF1p (ab216324) by LC-MS/MS were compared to IP IgG. For each condition (IP IgG, IP ORF1p) 5 independent immunoprecipitations were performed. Proteins with at least 2 distinct proteotypic peptides in all 5 replicates in the IP ORF1p condition with a ratio IP ORF1p/IP IgG > 2 and a p-value < 0.05 were considered as ORF1p interacting proteins. In the right panel, partners of ORF1p, which were not detected in the IP IgG condition (and therefore have no fold-change or p-value) are shown according to the number of peptides per 100 amino acids identified. **C.** Specific protein partners of ORF1p identified by proteomic mass spectrometry in LUHMES neurons, presented according to their fold change (ratio) and adjusted p-value. For each condition (IP IgG, IP ORF1p) 5 independent immunoprecipitations were performed. Proteins identified in the IP ORF1p (ab49) by LC-MS/MS were compared to IP IgG. Proteins with at least 1 distinct proteotypic peptide in at least 3 replicates identified in the IP ORF1p condition with a fold change > 2 and p-value < 0.05 were considered as ORF1p interacting proteins. In the right panel, partners of ORF1p, which were not detected in the IP IgG condition (and therefore have no fold-change or p-value) are shown according to their number of peptides per 100 amino acids identified. **D.** Top common GO categories (CC, MF, BP) of ORF1p protein partners in mouse (MN9D) and human (LUHMES) dopaminergic neurons **E.** Physical protein-protein interactions (PPI) network of ORF1p protein partners identified by mass spectrometry in MN9D neurons using the STRING database and visualized with Cytoscape. ORF1p protein partners belong to different functional and enriched GO terms (rings) : nucleus (dark blue), negative regulation of gene expression (blue), RNA polymerase II, core complex (light blue), synapse (red), neuron projection (pink) and RNA binding (green). Interactors previously identified are highlighted in yellow nodes. The thickness of the edges is proportional to the confidence in the physical interaction. Only proteins with physical interactions are shown. **F.** Physical protein-protein interactions (PPI) network of ORF1p partners found in LUHMES neurons by mass spectrometry using the STRING database and visualised via Cytoscape. Partners belong to different functional enriched GO terms (rings): gene expression (dark blue), nucleus (blue), axon guidance (red), mRNA splicing (light blue) and RNA binding (green). Interactors previously identified are highlighted in yellow nodes. The thickness of the edges is proportional to the confidence in the physical interaction. Only proteins with physical interactions are shown.

### ORF1p is located in the nucleus and associated to chromatin in post-mitotic neurons

Based on the identification of chromatin-related proteins as ORF1p protein partners, we characterized the subcellular localization of ORF1p in mouse and human neurons. Using immunofluorescence, we observed a predominant cytoplasmic expression of ORF1p in both, LUHMES and MN9D neurons, but also distinct nuclear dots in both species (Fig. 2A). Cytoplasmic and nuclear fractionation confirmed the presence of ORF1p in the nuclear fraction of mouse and human post-mitotic neurons (Fig. 2B). To gain a more precise understanding of the subcellular localization of ORF1p, we performed a more detailed biochemical fractionation, which confirmed that ORF1p is predominantly located to the cytoplasm, but also present in the membrane-bound fraction (Fig. 2C-E). Furthermore, ORF1p was localized to the soluble nuclear fraction (=nucleoplasm) and, importantly, reliably detected as chromatin-bound in the chromatin fraction of mouse neurons (Fig. 2C), human neurons (Fig. 2D) and in human post-mortem tissue extracts from the cingulate gyrus of a neurologically healthy individual (Fig. 2E). We confirmed the separation of all fractions using dedicated marker proteins and verified that stress granules were not contaminating the chromatin-bound fraction using the stress granule marker G3BP1 (Fig. 2C, D). The presence of ORF1p on chromatin is in line with previous observations indicating the association of ORF1p with chromatin independently of ORF2p in cancer cells^43^. Together, our observations suggest a close proximity and potentially functional relation of ORF1p with chromatin in neurons.

**Figure 2:**
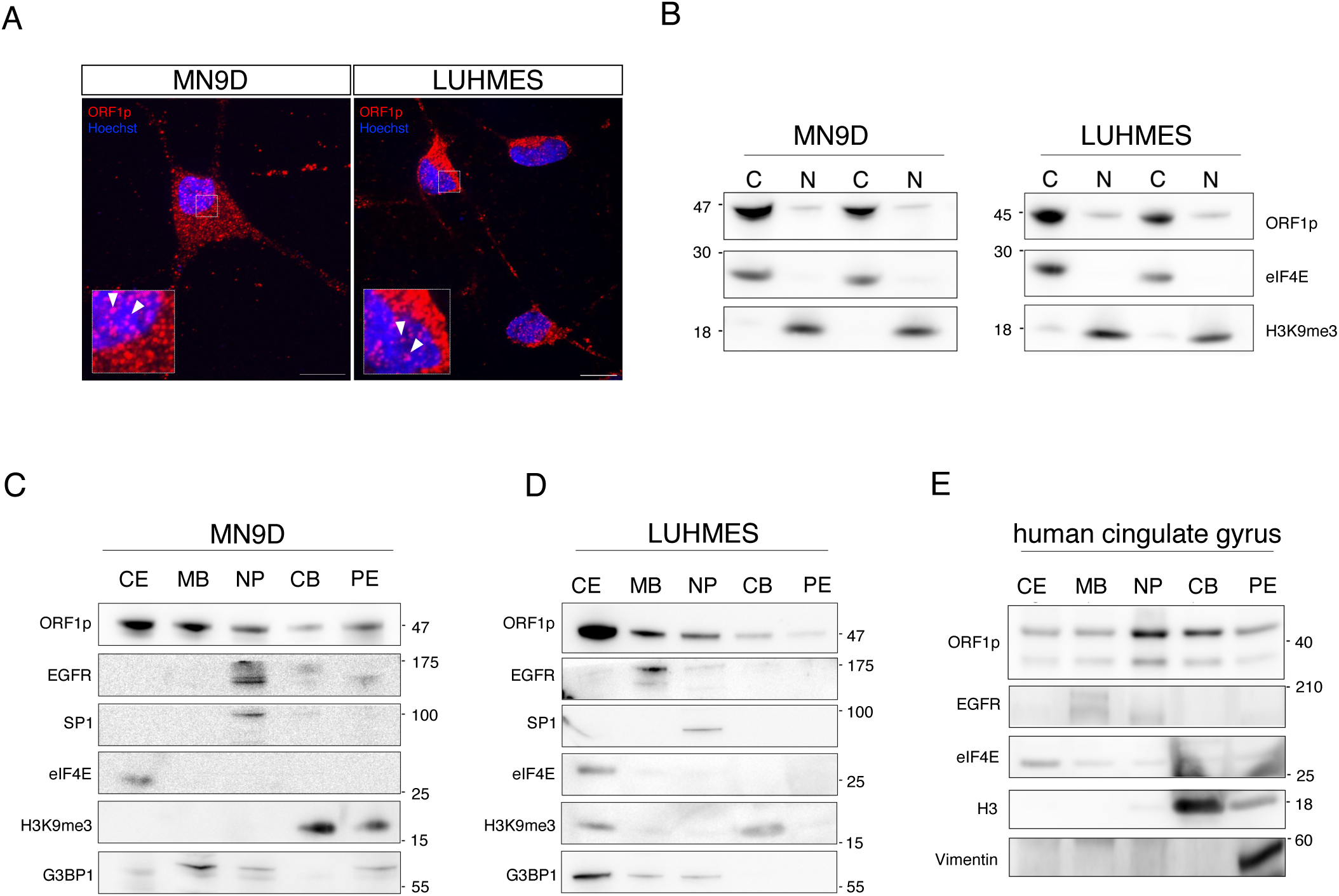
ORF1p is located to the nucleus and associated to chromatin in post-mitotic neurons. **A.** Immunostainings of endogenous ORF1p in LUHMES and MN9D neurons. Nuclear localisation of endogenous ORF1p is indicated by white arrows (antibody human ORF1p: Millipore MABC1152; antibody mouse ORF1p: Abcam ab216324) Scale bar is 10□µm. **B.** Biochemical separation of the nuclear and cytoplasmic compartments of two independent samples followed by Western blot using an ORF1p antibody and two markers, one for the nuclear compartment (H3K9me3) and one for the cytoplasmic compartment (eIF4E). ORF1p is present in the cytoplasmic and the nuclear fraction of LUHMES and MN9D neurons. A representative experiment is shown (n=3 independent experiments). *C: Cytoplasm, N: Nucleus*. **C.** Subcellular fractionation of the cytoplasm (CE), the membrane bound fraction (MB), the nucleoplasm (NP), the chromatin-bound fraction (CB) and the pellet (PE) of cell lysates of MN9D neurons followed by Western blotting using an ORF1p antibody and different protein markers (CE marker: eiF4E; MB marker: EGFR, NP marker: SP1; CB marker: H3K9me3; stress granule marker: G3BP1). ORF1p is present in all compartments, including the chromatin-bound fraction. A representative experiment is shown (n=3 independent experiments). **D.** Subcellular fractionation of the cytoplasm (CE), the membrane bound fraction (MB), the nucleoplasm (NP), the chromatin-bound fraction (CB) and the pellet (PE) of cell lysates of LUHMES neurons followed by Western blotting using an ORF1p antibody and different protein markers (CE marker: eiF4E; MB marker: EGFR, NP marker: SP1; CB marker: H3K9me3; stress granule marker: G3BP1). ORF1p is present in all compartments, including the chromatin-bound fraction. A representative experiment is shown (n=3 independent experiments). **E.** Subcellular fractionation of the cytoplasm (CE), the membrane bound fraction (MB), the nucleoplasm (NP), the chromatin-bound fraction (CB) and the pellet (PE) of lysates of human post-mortem cingulate gyrus tissue of a neurologically-healthy individual followed by Western blotting using an ORF1p antibody and different protein markers (CE marker: eiF4E; MB marker: EGFR, NP marker: SP1; CB marker: H3. PE marker: Vimentin1. ORF1p is present in all compartments, including the chromatin-bound fraction. A representative experiment is shown (n=3 independent experiments).

### Changes in nuclear ORF1p levels correlate with alterations in chromatin accessibility patterns in neurons

In order to explore whether ORF1p might have a functional impact on chromatin organization, we turned to an assay for transposase-accessible chromatin followed by sequencing (ATAC-seq). As we had previously shown that ORF1p was present in neurons in the human cingulate gyrus^17^ and established its presence in the chromatin-bound fraction in this brain region (Fig. 2E), we isolated nuclei from frozen human post-mortem cingulate gyrus tissues of six individuals (n=3 neurologically-healthy individuals and n=3 individuals with Parkinson disease (PD), Suppl Table 2). We then sorted them as schematized in Figure 3A using two antibodies, the neuronal marker NeuN, separating neurons from non-neuronal nuclei, and ORF1p, separating nuclei with high levels of ORF1p (ORF1p-high; P3, Fig. 3B) and those with lower levels of ORF1p including those not expressing ORF1p (ORF1p-low; P4, Fig. 3B). The corresponding isotype control profiles are shown in Suppl. Fig 2A. Confirming our previous observations of a predominant neuronal expression of ORF1p in the mouse and human brain^17^, non-neuronal nuclei were ORF1p negative, while neurons contained varying levels of nuclear ORF1p, allowing us to separate three distinct populations, NeuN-/ORF1p-minus; NeuN+/ORF1p-high and NeuN+/ORF1p-low, which we also verified by Western blot (Fig. 3C). This confirmed the absence of ORF1p in non-neuronal nuclei and the successful sorting of two neuronal populations with either high or low levels of nuclear ORF1p. We then performed high-throughput chromatin profiling on those three populations. In order to exclude that ORF1p-high and ORF1p-low nuclei were assimilated to a specific neuronal subtype, we plotted total ATAC-seq reads on promoter regions (<1kb) for cell type specific markers selected from Single Cell Base (https://biomarkerres.biomedcentral.com/articles/10.1186/s40364-023-00523-3) which showed a clear separation of NeuN- and NeuN+ accessibility profiles as expected and, importantly, no overt difference between accessibility profiles on neuronal subpopulation marker genes between ORF1p-high and -low nuclei (Suppl. Fig. 2B). Based on the number of normalized peaks, we determined differentially accessible regions (DARs). The Volcano plot in Figure 3D depicts all significantly enriched DARs, with 406 DARs more accessible in the NeuN+/ORF1p-high population (DAR+), and 379 DARs less accessible in NeuN+/ORF1p-high nuclei (DAR-≈ more accessible in NeuN+/ORF1-low neurons; log2 fold-change□□≥□1 / ≤ −1, p-value□≤ □0.01). We next analysed genomic features associated with each DAR. DARs were mostly located in introns (first and others) as well as in distal intergenic and, to a lesser extent, in promoter regions (Fig. 3E). However, there were significant differences in the distribution of DARs over genomic features between NeuN+/ORF1p-high and NeuN+/ORF1p-low nuclei. NeuN+/ORF1p-high/DAR− regions were enriched in introns (q = 0.0004; Fisher’s Exact test followed by Benjamini, Krieger and Yekutieli post-hoc test) compared to NeuN+/ORF1p-high/DAR+ regions, which were more frequently located in distal intergenic regions (q = 0.0004; Fisher’s Exact test followed by Benjamini, Krieger and Yekutieli post-hoc test). To gain a more precise idea about the functional features that DARs in both conditions were associated to, we overlapped DARs with regulatory features stemming from a ChromHHM annotation, which defines and characterizes chromatin states across the genome based on experimental data of a given tissue, in this case of human *post-mortem*cingulate gyrus (https://pmc.ncbi.nlm.nih.gov/articles/PMC8734071/; Fig. 3F). Less accessible chromatin (DAR−) in ORF1p-high nuclei coincided with enhancers (Enh), weak repressed polycomb (ReprPCWk) domains and domains associated with weak transcription (TxWk), while more accessible chromatin (DAR+) in ORF1p-high nuclei was mostly located in weak repressed polycomb (ReprPCWk) and weak transcription (TxWk) domains (Fig. 3F). The distribution of DARs between ORF1p-high and ORF1p-low nuclei was significantly different, with an overrepresentation of enhancers (Enh, q=0.0001) and active transcriptional start sites (TssA, q=0.0013) among DAR− regions and an overrepresentation of repressed polycomb regions (ReprPCWk, q=0.0001) and heterochromatin domains (Het, q=0.001) in DAR+ regions in ORF1p-high nuclei (Fisher’s Exact test followed by Benjamini, Krieger and Yekutieli post-hoc test). This indicated the existence of two different profiles of chromatin accessibility between these two nuclei populations. ORF1p-low nuclei were characterized by more accessible chromatin in intronic regions related to active transcription and enhancer activity and ORF1p-high nuclei showed open chromatin in repressed or heterochromatin regions rather located in distal intergenic genomic regions. We next visualized the top two candidates of each category using IGV. Tracks of DLX1 (distal-less homeobox 1) show an example of DAR+ regions in ORF1p-low neurons (Fig. 3G). DLX1 is a member of the homeobox transcription factor family, predicted to be a transcriptional regulator for TGF beta signalling and linked to autism^44^. The DAR in the DLX1 gene is annotated as a “flanking bivalent transcriptional start site/enhancer” and “bivalent enhancer” in adult cingulate gyrus ChromHMM annotations and overlaps with a CpG island. There are two other DARs in this region, one intergenic in-between DLX1 and DLX2 and one in the promoter region of DLX2, both characterized by bivalent enhancer or bivalent TSS annotations. These loci thus provide examples of the overrepresentation of DARs in ORF1p-low chromatin in intronic and promoter regions and transcriptional start sites and enhancers as depicted and quantified in Figure 3F. IGV tracks of SPACA3 (Sperm Acrosome Associated 3) show an example of a DAR+ region in ORF1p-high neurons (Fig. 3H).

**Figure 3:**
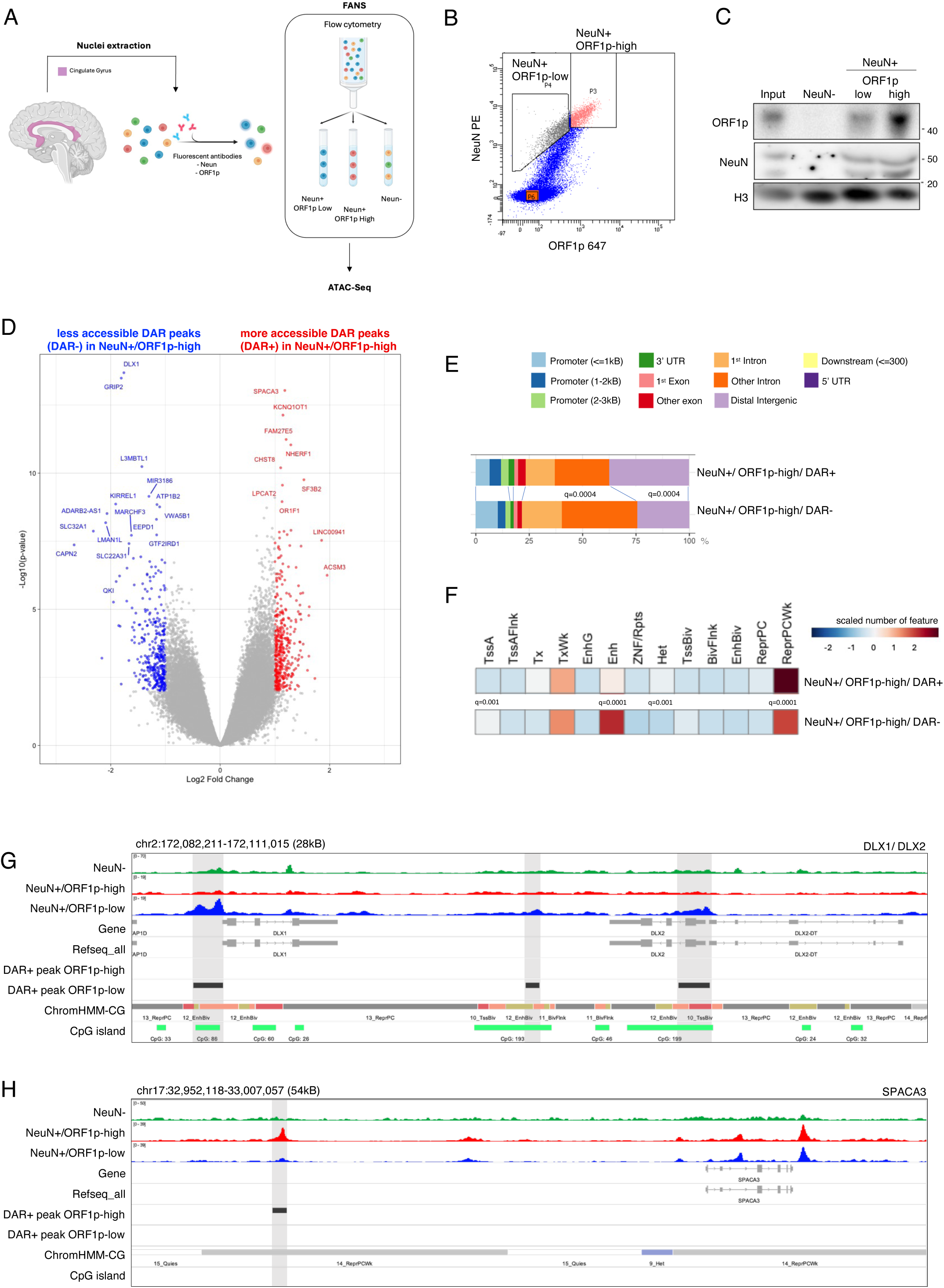
Nuclear ORF1p levels are associated with chromatin landscape variations in neurons. **A.** Graphical representation of the experimental set-up using fluorescent-activated nuclei sorting (FANS) to isolate neuronal nuclei with high or low nuclear ORF1p content from human post-mortem cingulate gyrus. Nuclei were extracted from pulverized frozen tissues of post-mortem cingulate gyrus of neurologically-healthy individuals (n=3) and individuals diagnosed with Parkinson’s disease (n=3). Nuclei were sorted with an antibody against NeuN to separate neurons (NeuN+) from non-neuronal cells (NeuN-) and an antibody against ORF1p to separate neuronal nuclei into two populations: those containing high amounts of ORF1p (ORF1p-high) and those containing no or low amounts of ORF1p (ORF1p-low). Sorted populations were then subjected to ATAC-sequencing. **B.** Representative fluorescent-activated nuclei sorting (FANS) profile showing the separation of non-neuronal cells (NeuN-; P6), NeuN+ cells (P3+P4) with either low (P4) or high (P3) nuclear ORF1p levels. **C.** Western blot of nuclei sorted by FANS. ORF1p is not detectable in non-neuronal cells (NeuN-). Higher amounts of ORF1p are present in the sorted population of ORF1p-high neuronal nuclei compared to ORF1p-low or ORF1p-negative neuronal nuclei. NeuN was used as a control for the separation of neuronal from non-neuronal nuclei. H3 was used as a loading control. **D.** Volcano-plot of differentially accessible regions (DARs) identified by ATAC-seq comparing ORF1p-high and ORF1p-low human post-mortem neurons LUHMES. Red: significantly more accessible regions (DAR+) in NeuN+ ORF1p-high; Blue: significantly less accessible regions in NeuN+ ORF1p -high (DAR−) (log2 fold change ≤ −1, pvalue ≤ 0.01). **E.** Genomic features associated with differentially-accessible-region (DAR) in NeuN+/ORF1p-high and NeuN+/ORF1p-low. **F.** Annotation of functional genomic features overlapping significant DARs using human post-mortem cingulate gyrus ChromHMM data retrieved from Roadmap (https://pmc.ncbi.nlm.nih.gov/articles/PMC873). **G.** IGV track of an example of DAR+ regions in ORF1p-low neuronal nuclei in the DLX1/2 genomic locus. **H.** IGV track of an example of a DAR+ region in ORF1p-high neuronal nuclei in the SPACA3 genomic locus.

SPACA3 is a sperm-specific gene and the DAR+ peak in ORF1p-high chromatin overlaps with a “weak repressed polycomb” annotation reflects the overrepresentation of ORF1p-high DAR+ regions in repressive chromatin regions (Fig. 3F). Using ChIP-seq data from sorted human post-mortem neurons^45^ for tri-methylated Histone H3 at lysine 4 (H3K4me3), which marks transcriptional start sites of active genes, tri-methylated Histone H3 at lysine 27 (H3K27me3), a histone modification associated with polycomb-mediated heterochromatin regions and acetylated Histone H3 at lysine 27 (H3K27ac), a marker for active enhancers, we observed significant differences in proportions of DAR+ regions overlapping with all three histone profiles. H3K27me3 was overrepresented in ORF1p-high nuclei (q=0.0003) in line with the repressive polycomb signature observed in the ChromHMM data, whereas H3K4me3 (q=0.0007) and H3K27ac (q=0.0003) were significantly overrepresented over DAR+ regions in ORF1p-low nuclei (Suppl. Fig. 2C, Fisher’s Exact test followed by Benjamini, Krieger and Yekutieli post-hoc test), in line with the active transcription and enhancer profile observed in the ChromHMM data (Fig. 3F). Together, this data indicated a distinct genomic and functional genomic profile of DARs dependent on ORF1p nuclear content, suggesting a regulatory function of nuclear ORF1p on chromatin dynamics. We then asked which transcription factor (TF) binding sites were overrepresented among the identified DARs and performed transcription factor motif enrichment analysis with HOMER (Suppl. Fig. 2D, E). In DAR+ regions in neuronal nuclei with a high ORF1p content, predicted TF motifs were predominantly showing members of the highly conserved basic leucine zipper (bZIP) transcription factor family like ATF3, involved in axonal regeneration^46^ and three members of the FOS TF family (FOSL2, FRA1 and FOS) which are TFs orchestrating the conversion of short-term extracellular signals into long-term functional changes via the activation of dedicated gene expression programs as members of the immediate early genes including roles in memory formation^47^. We also identified signatures of another immediate early gene, the zinc finger TF EGR1, within DAR+ regions in NeuN+/ORF1p-high nuclei. Interestingly, EGR1 is also involved in memory formation^47^. TF signatures on ORF1p-low DAR+ regions were mainly composed of homeobox genes like Engrailed-1 (En1) (Suppl. Fig. 2E) which have important functions in brain patterning^48^ but also in the maintenance of adult neurons^48^.

### LINE-1 knock-down induces changes in chromatin organization

Having established significant changes in chromatin accessibility in nuclei differing in the levels of ORF1p content in the human cingulate gyrus, we then turned to human LUHMES neurons in which we experimentally manipulated ORF1p steady-state levels using LINE-1 siRNAs targeting evolutionary young LINE-1 elements. LUHMES neurons were transfected at day three and retrieved at day seven of differentiation. Efficient knock-down of ORF1p was verified by immunostaining with a significant reduction in the cytoplasmic and the nuclear compartment (Fig. 4A) and by Western blot (Suppl. Fig. 3A) while LINE-1 RNA was slightly but not significantly reduced (Suppl. Fig. 3B). We also verified by Western blot the loss-of-function (LOF) of ORF1p after LINE-1 knock-down (KD) in the chromatin-bound fraction (Suppl. Fig. 3C). Using this knock-down protocol, we performed ATAC-seq on LINE-1 siRNA and control siRNA (Ctrl) transfected neurons. The MA-plot in Figure 4B shows the DARs (log2 fold-change□ ≥ □1 / ≤ −1, p-value□≤ 0.05), with 432 regions significantly more accessible (DAR+) after ORF1p-LOF and 917 less accessible regions (DAR−) compared to control. Similarly to human post-mortem nuclei with low ORF1p levels, DAR+ regions in the ORF1p-LOF condition were mostly located in introns and DAR− regions in distal intergenic regions, but with no significant overrepresentation when comparing the two conditions (Fig. 4C). When analysing the regulatory regions overlapping DAR+/DAR− regions in ORF1p-LOF nuclei (Fig. 4D) using a ChromHMM collection from human *substantia nigra* as a suitable reference for differentiated LUHMES cells which are dopaminergic^49^, we identified a very similar profile as in ORF1p-high and ORF1p-low nuclei in the cingulate gyrus (Fig. 3E). Indeed, just as in ORF1p-low nuclei, DAR+ regions in ORF1p-LOF nuclei were enriched on active transcriptional start sites (q=0.0001) and enhancers (q=0.03, Fisher’s Exact test followed by Benjamini, Krieger and Yekutieli post-hoc test). And similar to ORF1p-high nuclei, ORF1p-Ctrl DAR+ regions were overrepresented on repressed chromatin domains associated with polycomb (ReprPCWk, q=0.0001; ReprPC, q=0.005) and zinc finger genes/repeat associated genomic regions (ZNF/Rpts, q=0.001; Fisher’s Exact test followed by Benjamini, Krieger and Yekutieli post-hoc test, Fig. 4D). Again, using histone ChIP-seq data from sorted post-mortem neurons pooled from 5 different brain regions^45^, we also found a significant overrepresentation in the fraction of DAR+ regions in ORF1p-LOF nuclei over H3K4me3 domains (q=0.0001), while DAR− regions were overrepresented in H3K27me3 (q=0.0001) domains (Suppl. Fig. 3D). We then chose representative examples for DAR+ regions in ORF1p-Ctrl (Fig. 4E) and ORF1p-LOF (Fig. 4F) for visualization. WDR74 is an example of a DAR+ region in ORF1p-LOF neurons and is located in the promoter region overlapping with an active transcription start site (TssA) in line with the preferential location of DARs in ORF1p-LOF promoters depicted in Figure 4C and the overrepresentation of TssA regulatory regions more accessible in ORF1p-LOF (Fig. 4D). ROCK1P1, a pseudogene, contains two DARs overlapping with “ZNF/Rpts”, “Het” and “TxWk” annotations in line with regions overrepresented in ORF1p-LOF DAR− regions (Fig 4D). Next, we investigated the transcription factor binding sites enriched in those DAR. In DAR+ regions in Ctrl neurons, the family of basic helix–loop–helix (bHLH) TFs (Suppl. Fig. 3E) was strongly represented. These TFs have essential regulatory functions for neural cell fate specification and in embryonic development^50,51^. Interestingly, analogous to the TF profile overrepresented on DAR+ regions in ORF1p-low nuclei (Suppl. Fig. 2F), DAR+ regions in ORF1p-LOF nuclei were enriched in homeobox genes (Suppl. Fig. 3F). Therefore, low ORF1p nuclear content might favour accessibility to genomic regions regulated by homeobox proteins which are essential for specifying cell identities and positioning during embryonic development but also play a role in the adult brain related to neuronal maintenance^52^. Taken together, when comparing features across these two independent ATAC-seq experiments in human neurons (post-mortem cingulate gyrus neurons and differentiated dopaminergic neurons in culture) with high or low nuclear ORF1p content, we made similar observations; nuclei containing high levels of ORF1p showed more accessible chromatin preferentially in distal intergenic regions associated with polycomb-associated repressive domains and nuclei with low ORF1p content were predominantly characterized by more accessible DARs in intronic regions associated with enhancer functions and active transcription containing signatures of homeobox transcription factor binding sites (Suppl Table 3). This might suggest, that LINE-1 regulate cell identity in post-mitotic neurons.

**Figure 4:**
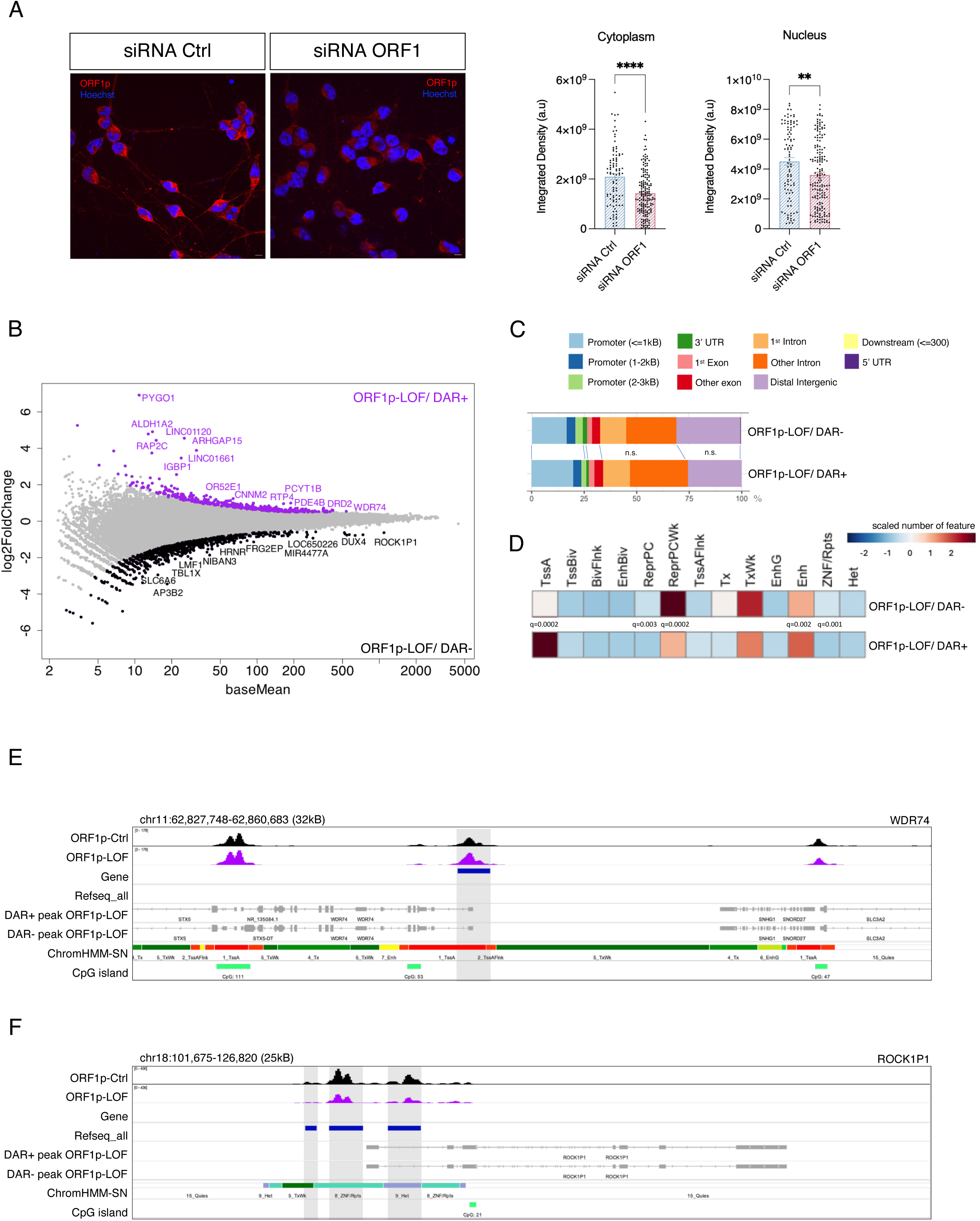
ORF1p knock-down induces changes in chromatin organization. **A.** Representative image of ORF1p knock-down by immunofluorescence (left) and quantification of ORF1p staining intensity (right; each dot on the dot blot represents one image). Two-tailed Mann–Whitney *U test*. Error bars are represented as mean ±□SEM. ∗p < 0.05; ∗∗p < 0.01; ∗∗∗p < 0.001; ∗∗∗∗p < 0.0001. **B.** MA-plot of differentially accessible regions (DAR) identified by ATAC-seq after ORF1p-LOF in human neurons (log2 fold change ≤ −0.5, p-value ≤ 0.05). **C.** Genomic features associated with differentially-accessible-regions (DARs) more (DAR+) or less accessible (DAR−) in ORF1p-LOF neurons. **D.** Annotation of functional genomic features overlapping significant DARs using human post-mortem *substantia nigra* ChromHMM data retrieved from Roadmap (https://pmc.ncbi.nlm.nih.gov/articles/PMC873). **E.** IGV track of a DAR+ region in ORF1p-LOF neuronal nuclei as identified by ATAC-seq in the WDR74 genomic locus. **F.** IGV track of DAR− regions in neurons after ORF1p-LOF in the ROCK1P1 gene locus.

### Nuclear ORF1p modulates chromatin accessibility on genes related to neuron-specific processes

We next compared the actual overlap of DAR-associated genes between both experimental systems. Of the 648 DAR-associated genes identified in cingulate gyrus neurons and the 1,189 DAR-associated genes identified in LUHMES neurons, 71 events were found to be shared between the two (Fig. 5A). This overlap represents 10.96% of the CG DARs and 5.97% of the LUHMES DARs. While this is only a modest overlap (Jaccard similarity coefficient 4.02%), GO analysis of these genes revealed five significant biological processes: axonogenesis, CNS neuron development, axon guidance, neuron recognition and neuron projection guidance (Fig. 5B). The GOChord plot in Figure 5B shows the eight genes sharing these GO terms. IGV tracks of two of them highlighted the presence of DARs on two enhancers in introns of the zinc finger transcription factor GLI2 and on a putative alternative promoter of XR_007087219 adjacent 5’ to GLI2 (ATAC-hu and ATAC-lu DARs) and an exonic DAR in a putative alternative promoter in the ROBO1 gene (ATAC-hu DAR) along an intronic DAR (chr3:79052746-79057310; ATAC-lu DAR) (Fig. 5C). These findings reinforce the possibility of a role of nuclear ORF1p in neuron-specific functions.

**Figure 5:**
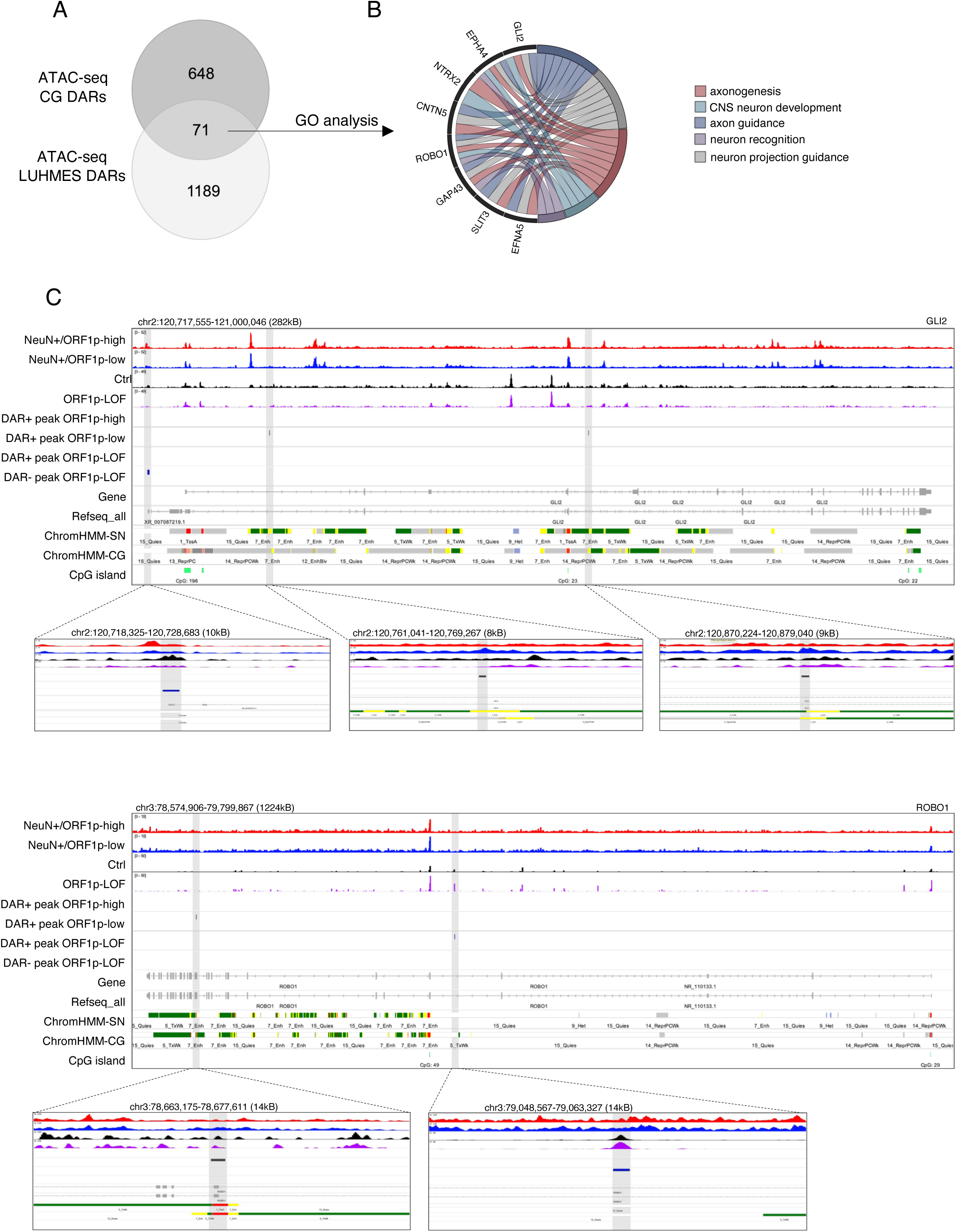
Nuclear ORF1p modulates chromatin accessibility on genes related to neuron-specific processes. **A.** Venn diagram showing the overlap between ATAC-seq DARs from human neuronal nuclei containing high or low amounts of ORF1p (1260 DARs) and ATAC-seq DARs from control human dopaminergic neurons (LUHMES) compared to neurons after ORF1p-LOF (719 DARs). 71 genes are associated to DARs in both conditions. **B.** GO analysis of 71 genes which are associated with at least one DAR in both ATAC-seq analyses. GO biological process complete; Fisher’s Exact test with FDR correction. 52 out of 71 were unambiguously mapped. Axon guidance (fold enrichment 11.92, FDR: 0.02, 7 genes). **C.** IGV tracks of 2 genes identified in the GO analysis in Figure 5B sharing DARs between human cingulate gyrus ORF1p-high/-low and human neurons Ctrl/ORF1p-LOF. Shown are genomic loci around the genes GLI2 and ROBO1.

### Knock-down of ORF1p affects the neuronal transcriptome, including downregulation of long genes related to neuron-specific functions

A way to interrogate changes in neuronal cell identity and function is via the characterization of the transcriptome. Therefore, we performed RNA-seq analysis after ORF1p-LOF in differentiated LUHMES neurons. Principal component analysis (PCA) separated ORF1p-Ctrl and ORF1-LOF transcriptomes (Fig. 6A). Using DESeq2, we identified 962 differentially expressed genes (DEG) (log2FC□□≥□0.5 / ≤ −0.5, p-value ≤ 0.01), of which 264 were upregulated and the majority, 698 genes, were significantly downregulated as shown in the Volcano plot in Figure 6B. Characterization of major “gene biotypes” using Ensembl annotations (hg38) revealed a similar pattern between all LUHMES expressed transcripts (total=56861) and transcripts upregulated upon ORF1p-LOF (total=264) with almost equal proportions of protein-coding transcripts (34% ORF1p-Ctrl; 29% ORF1-LOF) and long non-coding transcripts (30% ORF1p-Ctrl; 34% ORF1-LOF). However, ORF1p-LOF downregulated transcripts showed a significant different repartitioning of transcripts. Of all downregulated transcripts, 73 % were protein-coding and only 13% lncRNA transcripts (Chi-square test, observed versus expected, DF=4, p>0.0001). Similarly, when evaluating the fraction of long (>100kB) versus short genes (<100kB) differentially regulated upon ORF1p-LOF, we found that downregulated genes, but not upregulated genes, ware particularly long (>/=100kB). Compared to the fraction of long genes among all expressed transcripts in differentiated LUHMES neurons (9,69% of a total of 56861 transcripts) or genes upregulated upon ORF1p-LOF (10,61% of a total of 264 transcripts), downregulated genes upon ORF1p-LOF belonged more than two-times more frequent to the category of >100kB genes (21,78% of a total of 698 transcripts) (Fig. 6C, D). This might indicate that LINE-1 is implicated in the regulation of long gene expression. Interestingly, long genes have been associated with neuron-specific functions^53,54^. GO analysis of upregulated genes after ORF1p-LOF identified “odorant binding” and olfactory receptor activity as a significantly enriched (Suppl. Fig. 4A). More interestingly, downregulated genes were enriched in highly significant GO terms related to synapse (FDR: 2,53e-011, FE: 2.32; GO CC), neuron projection morphogenesis (FDR: 0,000425, FE: 2,72; GO BP), nervous system development (FDR: 0,000275, FE: 1,73; GO: BP) and synapse organization (FDR: 0.0005, FE: 3,01; GO BP) (Fig. 6E). Protein-protein interaction (PPI) network analysis of significant downregulated genes upon ORF1p-LOF contained in the GO CC term “Synapse” (GO:0045202; 50 genes) using STRING with a stringent cutoff (log2 fold change ≤ −0.5 / ≤ −0.5, p-value ≤ 0.005) showed a highly interconnected protein interaction network (Fig. 6F). A large proportion of these proteins were implicated in pre- and post-synapse, axon guidance, regulation of neurotransmitter levels and synapse organization. In line with this, KEGG pathway analysis identified “axon guidance” as one of the most significant pathways enriched in downregulated transcripts (Suppl. Fig. 4B). We independently tested four downregulated genes with functions related to neurons by RT-qPCR (DLG5, NEGR1, EFNA5 and GABRB3) and validated DLG5, NEGR1 and EFNA5 to be significantly downregulated after ORF1p-LOF (Fig. 6G).

**Figure 6:**
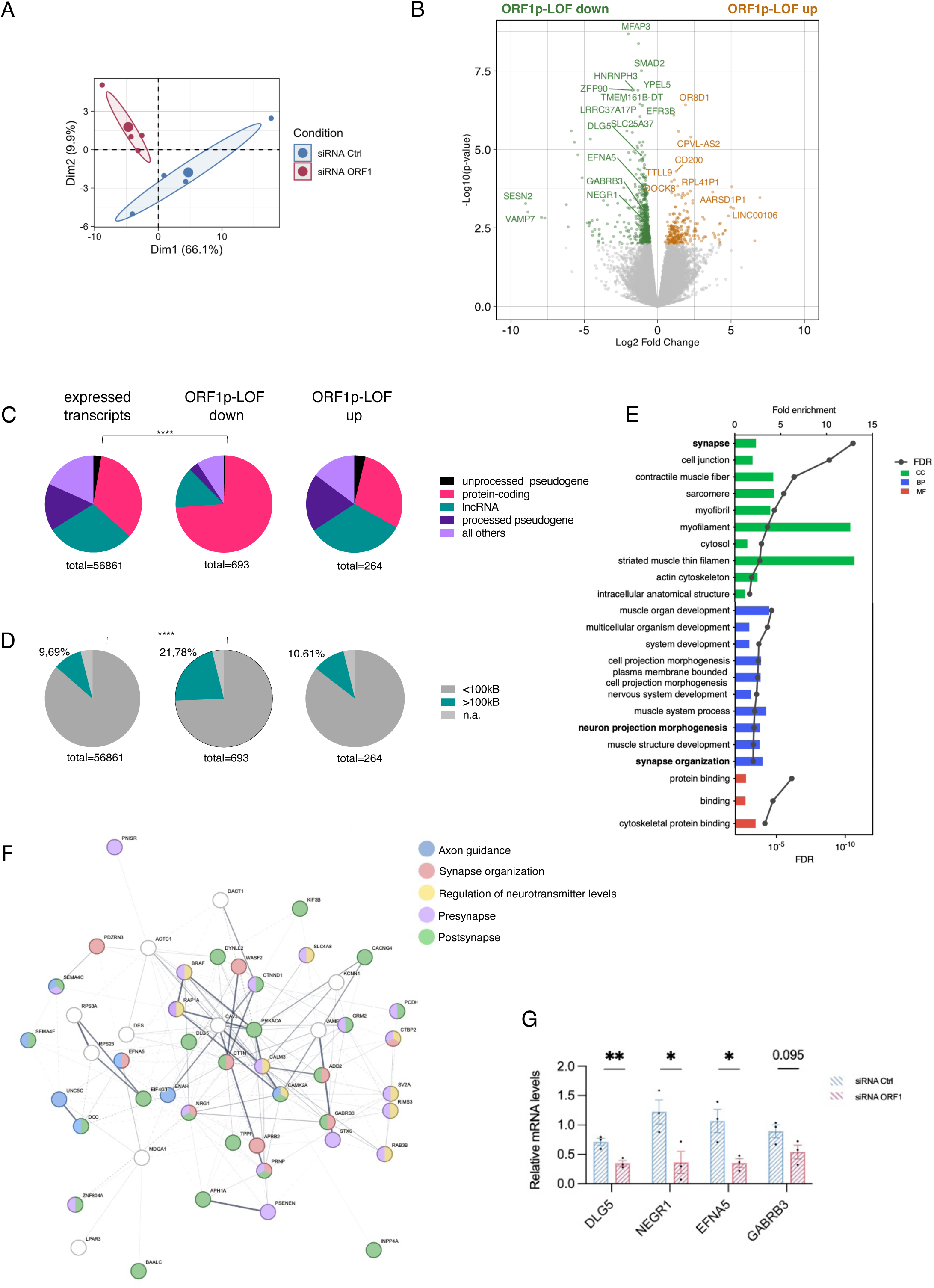
Loss of ORF1p leads to transcriptomic changes in neuron-specific genes. **A.** Principal component analysis (PCA) of RNA-seq data showing the separation of control neurons (blue) and ORF1p-KD neurons (red; n=4 samples per condition, LUHMES). **B.** Volcano plot showing differentially expressed genes (DEGs) in LUHMES neurons upon ORF1p-LOF. 698 genes were significantly upregulated (in red: log2 fold change ≥ 0.5, p-value□≤ 0.01); 264 genes were significantly downregulated (in blue: log2 fold change ≤ −0.5, p-value ≤ 0.05). **C.** Parts of whole graph showing the distribution of gene biotypes extracted from NCBI RefSeq among all expressed transcripts present in the RNA-seq analysis (left) or upregulated (middle) or downregulated transcripts (right) upon ORF1p-LOF. Distributions of gene biotypes among expressed or upregulated transcripts were similar while downregulated transcripts were mainly protein-coding transcripts. Gene biotype overrepresentation analysis was done using a Chi-Square test with observed = number of feature counts in ORF1p-LOF and expected = all RNA-seq transcript counts. **D.** Gene length analysis of all expressed transcripts in human LUHMES neurons (left), upregulated (middle) or downregulated transcripts (right) upon ORF1p-LOF. Genes were separated in two categories; long genes (>100kB) and short genes (</=100kB). The fraction of long genes among all and upregulated transcripts was similar (around 10%) but was significantly higher among downregulated transcripts (22%). Long gene overrepresentation analysis was done using a Chi-Square test with observed = number of feature counts in ORF1p-LOF and expected = all RNA-seq transcript counts. **E.** GO analysis of significant downregulated genes from ORF1-LOF vs Ctrl. FDR was calculated from p-values (Fisher’s exact test) by the Benjamini-Hochberg procedure. CC: Cellular Component, BP: Biological process and MF: Molecular Function. **F.** Protein-protein interaction (PPI) network of significantly downregulated genes upon ORF1p-LOF (log2 fold change ≤ −0.5, p-value ≤ 0.005) illustrated for the GO Cellular Component term “Synapse” GO:0045202 (50 genes). Genes are coloured according to their associated GO term. **G.** Quantitative RT-PCR depicting the relative mRNA expression of DLG1, NEGR1, EFNA5 and GABRB3 normalized to ENO2, TBP and HPRT in LUHMES neurons treated with siRNA Ctrl or siRNA ORF1 (n□=□3 per condition). Two-tailed Student’s t-test. Data is represented as mean□± SEM ∗p < 0.05; ∗∗p < 0.01; ∗∗∗p < 0.001; ∗∗∗∗p < 0.0001.

DARs identified in ATAC-seq are associated to the closest gene, although this might not be always biologically relevant as open chromatin regions, especially outside proximal promoter regions, do not necessarily regulate proximal genes but might act at greater distances. We nevertheless overlapped RNA-seq DEGs and ATAC-seq DARs obtained from LUHMES neurons and identified an overlap of 26 genes (Suppl. Fig. 4C). Interestingly, GO analysis of these genes (APBB2, CACNA2D1, EVC, DUSP22, ENOX2 TMEM120B, ATF7, NMNAT2, USP13, SEPTIN9, SPATS2L, MDGA1, SMAD2, EFNA5, PSMB11, CPVL-AS2, UXS1, PPM1L, PCDH8, SDC3, PTBP2, PTPN3, MARCHF1, AUTS2, SATB1, TMTC2) revealed that they belonged to neuron specific pathways such as “axon development”, “regulation of synaptic membrane adhesion” and “neuron migration” (Suppl. Fig. 4C). Interestingly, three of these genes with neuronal functions, namely AUTS2, PTBP2 and EFNA5, also contained DARs in cingulate gyrus neurons. AUTS2 (Activator Of Transcription And Developmental Regulator) is a component of the polycomb group multiprotein complex 1 with regulatory functions in neurite morphology and is associated with autism spectrum disorders, intellectual disability and developmental delay^55^. PTBP2 (Polypyrimidine Tract Binding Protein 2) is predominantly present in the brain and involved in splicing regulation^56^. EFNA5 (Ephrin-A5) is involved in migration, repulsion and adhesion, notably in axon guidance during neuronal development^57^. The gene locus of EFNA5 with ATAC-seq and RNA-seq tracks is shown in Suppl. Figure 4D.

Taken together, these transcriptomic changes upon ORF1p-LOF suggest a role of steady-state ORF1p in the homeostasis and maintenance of post-mitotic neurons.

### ORF1p loss of function leads to changes in neurites morphology

In addition to the fact that nuclear ORF1p dependent changes in chromatin accessibility and in the transcriptome converged upon the notion that ORF1p might be involved in regulating the homeostasis and maintenance of adult neurons at steady-state, the mass spectrometry analysis of protein interactors of endogenous ORF1p in both, mouse and human neurons, had revealed proteins related to neuron-specific functions with high-confidence (cat.II ORF1p interacting proteins, Fig. 1D-F). Cat.II ORF1p interactors belonged to GO categories like “neuron projections” (33 proteins in LUHMES, 65 in MN9D; 5 in common), “synapse” (42 proteins in LUHMES, 111 in MN9D, 15 in common), “dendrites” (18 in LUHMES, 43 in MN9D, 5 in common) and “axo-dendritic transport” (6 in LUHMES, 8 in MN9D, 1 in common; Suppl Table 1). Among these cat.II protein interactors of endogenous mouse and human ORF1p in neurons, 18 common proteins emerged; 9 ribosomal proteins and 2 myosin proteins with non-canonical functions in addition to YBX1, HNRNPU, MLF2 and FXR1 and two microtubule-associated proteins with canonical functions in neurite morphology, namely two microtubule-associated proteins, MAP6 (dendrite morphogenesis, axonal growth and transport^58^, and MAP1B (neurite extension^59^). These findings again suggest a possible role of ORF1p specific to neurons which converge towards a pathway related to neurites (Fig. 1D). Moreover, this idea was further supported by the presence of ORF1p itself in neurites as revealed by immunofluorescence stainings of endogenous ORF1p in MN9D and LUHMES (Fig. 7A). Using TUJ1 to identify neurites, immunostaining of ORF1p showed its localization in neurites in both, human and mouse dopaminergic neurons (Fig. 7A). Additionally, we transfected MN9D neurons with human ORF1p-HA and detected ORF1p by HA immunostaining, as expected in the cytoplasm and the nucleus, but also localized to neuronal extensions (counterstained with TUJ1), including axons (counterstained with NEFL) (Fig. 7B). This indicated a possible colocalization of ORF1p with proteins related to neurite functions in the neuritic compartment itself. We verified the co-localization and co-immunoprecipitation of PABPC1, an interesting candidate with regard to neurite physiology^60,61^ and protective against dopaminergic neurodegeneration^62^, which we had identified as interacting with ORF1p in both, human and mouse neurons by mass spectrometry (Suppl Table 1), and others in different cultured cell types^26–30,33,63^. Co-immunoprecipitation (Fig. 7C) and colocalization (Fig. 7D) in mouse neurons validated the interaction and identified that both proteins colocalize in the cytoplasm but also in neurites. We thus hypothesized, that both, nuclear ORF1p-induced chromatin and transcriptional changes and ORF1p protein-protein interactions in the cytoplasm and in neuronal extensions might affect neurite morphology. In order to test this hypothesis, we carried out ORF1p-LOF in MN9D neurons at seven days of differentiation and collected them at day 10 to perform neurite staining. ORF1p-LOF in mouse neurons was efficient as verified by Western blot (Fig. 7E). Using an automated, non-biased image quantification workflow, we measured neurite morphology via measures of neurite diameter and area (Fig. 7F, G). Quantifications of images illustrated in Figure 7F revealed a significant loss of the mean diameter of neurites after ORF1p-LOF accompanied by a decrease in the total neurite area (Fig. 7G). In summary, these data suggest that ORF1p plays a role in the maintenance of neurite morphology possibly by converging pathways mediated by protein interactions and by modulating chromatin accessibility and transcriptomic landscapes.

**Figure 7:**
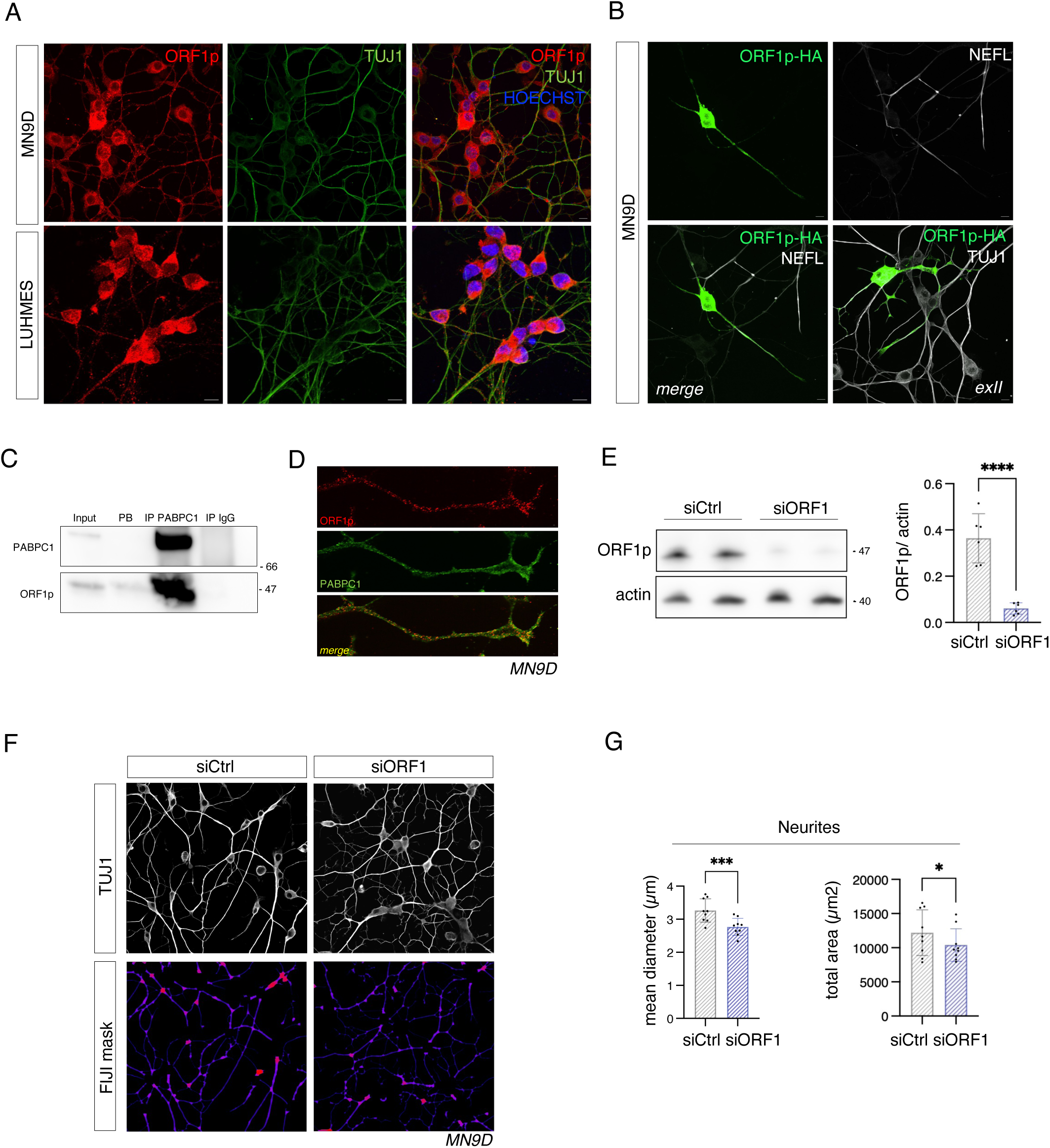
ORF1p knock-down induces phenotypic changes of neurites. **A.** Representative images showing ORF1p expression (red) in neurites (green; TUJ1) in mouse (upper panel) and human (lower panel) dopaminergic neurons. Scale bar is 10□µm. **B.** Expression of plasmid encoded human ORF1p tagged with HA and labelled with a HA antibody (green) in differentiated mouse dopaminergic neurons (MN9D). Neurons were transfected at day 10 of differentiation. Neurites were counterstained with NEFL or TUJ1 (white) as indicated. ORF1p is readily detectable in TUJ1-positive neurites and NEFL-positive axons. Scale bar 10□µm. **C.** Co-immunoprecipitation of ORF1p with PABPC1-antibody coated beads. Western blot confirming efficient immunoprecipitation of PBPC1 and specific co-immunoprecipitation of ORF1p. **D.** Immunofluorescence staining of ORF1p (red) and PABPC1 (green) in differentiated mouse neurons. Co-localization is visible in yellow. **E.** Western blot analysis and quantification of ORF1p knock-down in differentiated MN9D (n = 6, one dot in the dot plot corresponds to one well; Mann Whitney test, ∗∗∗∗p < 0.0001) (n=3 independent experiments). **F.** Immunofluorescent staining of the neuritic network of mouse dopaminergic neurons using TUJ1. siRNA Ctrl neurons are shown on the left and siRNA ORF1p neurons on the right. Scale bar 10□µm. Masks were established using a Fiji-operated script for the quantification of neurite morphology. **G.** Quantifications of neurite morphology based on TUJ1 staining of neurites. The mean diameter of neurites (in µm) was quantified using Fiji. Each point on the graph corresponds to the mean diameter value of an independent experiment. Neurite total area (µm2) was quantified in siRNA Ctrl and siRNA-ORF1p MN9D neurons; n=9 independent experiments.

## Discussion

Starting from the surprising observation that the LINE-1 encoded ORF1p protein is expressed at steady-state in various neuronal subtypes in the adult brain of mice and humans^17,18^, we addressed here the hypothesis that the sustained presence of ORF1p in an important fraction of neuronal cells in the brain^17^ might be a requirement for these neurons to maintain specific functions. Using quantitative mass spectrometry, confocal imaging, chromatin accessibility and transcriptomics on human post-mortem neurons and differentiated mouse and human neurons in culture with varying levels of nuclear ORF1p, we provide evidence that ORF1p interacts with chromatin-binding proteins, binds to chromatin, modulates chromatin accessibility and maintains neuronal gene expression. Loss of function of ORF1p leads to the downregulation of genes related to neuronal processes and alters neurite morphology. In addition, ORF1p directly binds to proteins implicated in neurite morphology, indicating that it might be the converging interaction of ORF1p with proteins and chromatin that mediate this effect. To our knowledge, this data represents the first demonstration of a role of a flLINE-1 encoded protein in the homeostasis of mature neurons and indicates a possible exaptation of a retrotransposon-encoded protein for physiological functions in the adult brain. This data thus adds to the growing evidence for the exaptation of retrotransposon-derived RNA and proteins, for now documented during early development^2,10,11,64^, including in the developing brain^3^. These studies have focused on the role of LINE-1 RNA. During early preimplantation development, LINE-1 RNA transcription leads to global chromatin relaxation and ribosomal DNA transcription promoting blastocyst formation^10,11^. However, accumulating evidence also suggests that LINE-1, not necessarily full-length and coding, interfere in the differentiation of the neuronal lineage. LINE-1 RNA was identified as a structural organizer of chromatin organization^65^, activators of neuronal gene expression in ESCs^66^, in human neuronal progenitors^67^ and cerebral organoids^12,68^, as well as in embryonic and adult neurogenesis^3^ and as regulators of mouse brain corticogenesis^14^. Importantly, one study reported that downregulation of LINE-1 expression in *cis* using CRISPRi in cerebral organoids led to the upregulation of genes linked to neuronal growth and neurogenesis and resulted in reduced cerebral organoid size^68^. Another study using shRNA to knock-down LINE-1 in *trans* reported the downregulation of neuronal and synaptic genes^14^, similar to what we observed in post-mitotic neurons here. Mangoni et *al.* showed that LINE-1 act as a long non-coding (lnc) RNA and as such epigenetically mediates the deposition of H3K27me3 on these downregulated genes regulating cell differentiation and neuronal migration in the developing mouse brain. They did not investigate, however, whether during corticogenesis, ORF1p was present on chromatin and/or attached to the LINE-1 RNA. While this would be interesting to pursue, both sets of data suggest a function of LINE-1 and/or LINE-1 encoded ORF1p in regulating parts of the neuronal transcriptome, either during cortical development^14^, or, as we show here, in the maintenance of a stable post-mitotic phenotype. Further, both studies link LINE-1 activity, either mediated through the binding of the LINE-1 RNA to polycomb complexes^14^ or via nuclear ORF1p mediating differential accessibility on polycomb-repressed genome domains itself (this study: higher nuclear levels of ORF1p, both in human post-mortem and in LUHMES neurons, are associated with more accessible chromatin on weak polycomb repressive domains), by affecting chromatin accessibility on the PRC-related gene AUTS2 or by interacting with the PRC1 protein CBX8 (this study). PRC1 is involved in chromatin silencing following PRC2 mediated tri-methylation of histone H3 at lysine 27^69,70^. Notably, Polycomb complex proteins are important for cellular identity in development^71^ and in neuronal morphology^72^ through the epigenetic silencing of target genes^73^. LINE-1 RNA and PRC2 interact also outside the nervous system maintaining nucleolar organization and an embryonic stem cell identity^74^. The interaction of LINE-1 RNA with PRC2 or the differential accessibility of PRC chromatin domains depending on nuclear ORF1p content and the interaction of ORF1p with the PCR1 protein CBX8 could thus suggest a global and important role in maintaining cell identities, possibly including that of post-mitotic neurons. While the analysis identifies a clear association between ORF1p levels and chromatin accessibility patterns, other factors such as disease status may also contribute to the observed differences. However, the consistency between post-00mortem findings and controlled *in vitro* experiments suggests a robust relationship between ORF1p and chromatin accessibility independent of these potential confounders and provides cross-validation across different experimental systems, supporting the specificity of ORF1p’s effect on chromatin accessibility.

One important question which arises in this context is that of the molecular form of LINE-1 associated with the observed cellular and molecular changes upon knock-down, that is whether ORF1p is acting alone, ORF1p is associated to LINE-1 RNA, to other cellular RNAs and/or to ORF2p. As introduced above, ORF1p is a nucleic acid chaperone and a RNA binding protein and known to be, due to its *cis*-preference, associated to LINE-1 RNA but also to other cellular RNAs prone to forming a LINE-1 RNP^75–79^. We have shown here that ORF1p is present in the nucleus and bound to chromatin in post-mortem human neurons and in differentiated mature mouse and human neurons in culture, an observation that had been already made earlier in other cell types and *in vitro*^80–82^. In dividing cells, the LINE-1 RNP is supposed to enter the nucleus passively when the nuclear envelope breaks down during mitosis^35,82^ before ORF1p re-exits the nucleus through an active process^35^. In non-dividing somatic cells^38^ and in G_o_-arrested cells like neurons^37^, where retrotransposition events can occur, ORF1p supposedly enters the nucleus through binding to import proteins and active nuclear transport^41,83^ including in post-mitotic neurons^19^. Whether once in the nucleus, it associates with RNA or not to bind chromatin, is an open question, which remains to be addressed. It is however possible, that the presence of steady-state nuclear ORF1p could be a particular feature of post-mitotic neurons providing a unique nuclear function. As we have shown here, nuclear chromatin containing higher or lower levels of ORF1p showed differences in chromatin accessibility with similar characteristics in two different neuronal systems, suggesting a functional role of ORF1p on chromatin which might or might not be involving (LINE-1) RNA. While the overlap of genomic sites of differential chromatin accessibility in the cingulate gyrus and in LUHMES neurons was rather low, both systems recapitulated specific features of chromatin accessibility depending on nuclear ORF1p content. High nuclear ORF1p led to more accessible chromatin on weak repressed polycomb domains and predominantly in distal intergenic regions while low nuclear ORF1p was associated with more accessible chromatin over enhancers and flanking active transcription sites in intronic regions. How exactly nuclear ORF1p can regulate chromatin accessibility remains to be investigated. One possibility is through the association with proteins involved in chromatin-related functions. Indeed, we identified many nuclear proteins with chromatin-related function interacting with ORF1p. These protein interactors of ORF1p provide an extensive network of proteins related to chromatin organization and transcription regulation. Three protein interactors of ORF1p stood out as they were identified as common ORF1p interacting proteins in both, mouse and human neurons: RCOR2 (corepressor 2 of REST), a transcriptional repressor required for mouse ESC differentiation^84^ and neurogenesis^85,86^, YTHDC1, a protein implicated in m6A-dependent RNA metabolism^87^ and HNRNPU, a DNA and RNA binding protein associated with various neurodevelopmental disorders^88^. HNRNPU is involved in chromatin organization, transcription regulation and many other nuclear processes^89^ and has been previously identified as an ORF1p interactor in cancerous cell lines^33^. Interestingly, YTHDC1 has been shown to bind m6A-modified LINE-1 RNA in order to form a protein-RNA complex which represses the 2-cell program in ESCs^90^ but might also be involved in neuronal survival^91^ and cerebral stress response^92^. It would be interesting to test the functional relevance of the interaction of ORF1p with these proteins in mediating chromatin accessibility in neurons. While the functional impact of these protein-protein interactions remains to be determined, their identification reinforces the hypothesis that it might be ORF1p that modulates chromatin accessibility in mature neurons, either directly or indirectly through the binding to these proteins and/or to RNA. Given the extensive list of ORF1p protein interactors with functions in RNA metabolism, we cannot exclude that ORF1p might also regulate gene expression post-transcriptionally. While many of these interactions were consistent with previoulsy identified protein partners of ORF1p in human cancer cell lines, mouse brain tissues and sperm and were identified in well-established neuronal cell lines, future studies in primary neurons or brain tissue will be important to confirm their relevance *in vivo* and fully characterize the specificity and strength of these associations.

A combination of observations suggests a role of ORF1p in neuron-specific functions. (i) shared genes containing DARs in both neuronal systems were enriched in gene categories related to axonogenesis and neuron projection. (ii) Transcriptomic changes observed after ORF1p-LOF indicated that downregulated transcripts were particularly long, protein-coding and related to neuron-specific functions like synapse, neuron projection and axon guidance. (iii) Endogenous and HA-tagged ORF1p localized to neurites, both in mouse and human neurons and (iv) mass spectrometry identified axono-dendritic ORF1p interacting partners. (iv) Functionally, ORF1p-LOF induced morphological changes in neurites. Indeed, effects of LINE-1 on neurite morphology have been previously observed during mouse and human brain development. LINE-1 KD *in vitro* and *in vivo* induced morphological NPC differentiation and increased the number of dendritic branches, length and complexities^13^. This was accompanied by LINE-1-KD DEGs enriched in functional categories including morphogenesis and projection, similar to what we identified here. Morphological changes during neurodevelopment were characterized by an increase in neurite branches, while we observed decrease in neurite diameters. This suggest that flLINE-1 activity acts on neuronal morphology in post-mitotic neurons. Interestingly, while Toda *et al.* did detect LINE-1 RNA on chromatin, there was no correlation of the presence of RNA and DEG gene loci as described during zygotic development^10^, which could suggest that the presence of ORF1p is mandatory for specificity in chromatin targeting in neurons. Indeed, during pre-implantation development, ORF1p is exclusively cytoplasmic^10^, while in neurons, ORF1p is readily detectable in the nucleus at steady-state. This data suggests a dual role of ORF1p converging on the regulation of specialized neuronal functions. While future work will be required to demonstrate the direct causal links between these sequential events, the consistent enrichment for neuron-specific biological processes such as ‘axon guidance’ in the proteomic, chromatin, and transcriptomic analyses provides strong correlative support for a unified functional pathway.

Together with mounting evidence of the importance of LINE-1 activity during embryonic and adult neurogenesis^2^ and neurodevelopment^3^, our work thus contributes to the increasing evidence of the exaptation of what was once called “selfish elements” for important physiological brain functions and beyond. This is important to take into consideration when targeting LINE-1 activity for developing novel therapies for diseases like cancer, autoimmune and neurodegenerative diseases. We have shown previously, that excessive ORF1p and its translocation to the nucleus alters nuclear membrane integrity, transport and heterochromatin organization^19^, indicating that increased nuclear levels of ORF1p might be deleterious. Our present study adds to the understanding of LINE-1 biology by showing that mature neurons depend on ORF1p for the fine-tuning of chromatin organization, neuron-related gene expression and neurite morphology. Taken together, we propose a threshold model in which intermediate levels of ORF1p are necessary for maintaining neuronal morphology and function while levels going below or beyond a physiological steady-state expression window might be harmful to neurons.

## Material and Methods

### Ethical approvals

Reglementary authorizations to use human cell lines and human samples were obtained from the French Ministry of Higher Education, Research, and Innovation (authorization number DC-2020-4013). Experiments using siRNA in human cells (LUHMES) were authorized according to regulatory procedures defined by the French Ministry of Higher Education, Research, and Innovation (OGM n°8273 and OGM n°10463).

### Cell culture

Lund Human Mesencephalic (LUHMES) cells were ordered from ATCC (ATCC-CRL-2927). Cell culture dishes were pre-coated with 50 μg/mL poly-L-ornithine (Merck, # P4957) diluted (1:2) in DPBS (Gibco, # 14190144) with additionally 1 μg/mL human plasma fibronectin (Sigma, # F0895) at 37°C for at least 3 hours and then rinsed 2 times with DPBS. Proliferating LUHMES were cultured using Advanced DMEM/F12 (Gibco, #12634028) added with 1% N-2 supplement (Gibco, # 15410294), GlutaMax (Gibco, # 35050061) and 40 ng/mL human recombinant FGF (Peprotech, # 100-18B) at 37C° in 5 % CO_2_ incubator. LUHMES cells were differentiated into mature dopaminergic neurons. For this purpose, neurons were cultured in Advanced DMEM/F12 added with 1% N-2 supplement and GlutaMax, supplemented with 1 µg/ml doxycycline (Sigma), 2 ng/ml recombinant human GDNF (Peprotech, # 450-10) and 1mM cAMP (Sigma, # D0627). ORF1p immunoprecipitation was done on day 5 of differentiation and all ORF1p knock-down experiences were conducted on day 7 of differentiation, when LUHMES cells are mature dopaminergic neurons^93,94^. Medium was changed every two days. Cells used in this study had a maximum of 15 passages.

The MN9D cell line was purchased from Merck (# SCC281). Cell culture plasticware was coated with 1 mg/mL poly-lysine (Merck, # P4707) for 2 hours and then rinsed 2 times with DPBS. Cells were cultured in DMEM High Glucose (Sigma, # D5796) supplemented with 10% fetal bovine serum (FBS) (Gibco, #10270098) in 5 % CO_2_, 37°C incubator. MN9D were differentiated for 10 days with the following medium: DMEM High Glucose (Sigma, # D5796) supplemented with 10% fetal bovine serum (FBS) (Gibco, #10270098) with 1 mM n-butyrate (Sigma, # B5887) and 1 mM dibutyryl cAMP (Sigma, # D0627). Media was changed every two days. Experiences were performed on day 10 of differentiation. Cells used in this study had a maximum of 15 passages.

### SiRNA delivery

Around 150 000 cells/well of LUHMES or MN9D were plated. Media was freshly replaced 2 hours before. Cells were lipofected using RNAiMAX (Invitrogen, # 13778150) at day 3 of differentiation for LUHMES and day 7 of differentiation of MN9D, with 100 nM of siRNA.

siRNA Ctrl: TAATGTATTGGAACGCATA

siRNA Orf1^95^ (human ORF1 - LUHMES): AAGAAGGCTTCAGACGATCAA

siRNA Orf1^20^ (mouse ORF1 - MN9D): CTATTACTCTGATACCTAAAC

### Plasmid delivery

MN9D were plated at the density of 200 000 cells/well into a pre-coated 4-well plate. On day 7 of differentiation, new pre-warmed media was added 2 hours before and cells were transfected with 1 µg of plasmid/well (pcDNA3-ORF1-HA, kind gift from Dr. John and Dr. Ariumi^96^) or pmaxGFP (Lonza) using Lipofectamin 3000 (Thermo Fisher Scientific, # L3000015) according to manufacturer’s instruction.

### Western Blot

Cells were scraped at day 7 of differentiation (LUHMES) and at day 10 of differentiation (MN9D) in RIPA buffer (10mM Tris-HCl pH 8.0, 150mM NaCl, 1mM EDTA, 1% Triton X-100, 0.1% Sodium Deoxycholate, 0.1% SDS). Laemmli buffer was added, and samples were boiled 10 min at 95°C before loading on a 1.5mm NuPAGE 4-12% Bis-Tris polyacrylamide gel (Invitrogen, #NP0336BOX) for the Western blotting. Gels migrated in NuPAGE™ MES SDS Running Buffer (Invitrogen, #NP0002) for 1h10 minutes at 175mV. Gels were transferred onto a methanol-activated PVDF membrane (Immobilon, # IPVH00010) in transfer buffer (25 mM Tris pH:8,3 and 192 mM Glycine) during 1h30 at 400 mA. Membranes were blocked in 5% milk TBST (0,2% Tween 20, 150 mM NaCl, 10 mM Tris pH:8) for 1 hour. Primary antibodies were diluted in 5% milk TBST, and membranes were incubated o/n at 4°C. After 3 x 10 min washing in TBST, membranes were incubated for 1h with the respective secondary antibodies diluted at 1/2000 in 5% milk TBST, followed by 3 x 10 min washes in TBST. Membranes were revealed by the LAS-4000 Fujifilm system using Clarity Western ECL Substrate (Bio Rad) or Maxi Clarity Western ECL Substrate (Bio Rad) according to the sensitivity needed. Quantifications were performed using LI-COR Image Studio Lite (v.5.2.5).

### Immunofluorescence

Cells were fixed using 4% methanol-free formaldehyde (Sigma, # 28908) diluted in PBS for 20 min at RT and washed 3 times with PBS. Cells were permeabilized with 0.5% Triton X-100 PBS 20 min and washed twice with PBS. 100mM glycine was added for 20 min and after washing 3 times with PBS, cells were incubated with blocking buffer (10% FBS, 0.5% Triton X-100, PBS) for 1 hour. Primary antibodies in blocking buffer were added overnight at 4°C. The next day, following 3 washes with PBS for 10 min, cells were incubated with secondary antibodies diluted in blocking buffer for 1 hour at RT. Finishing with 3 PBS washes, slices were mounted with Fluoromount (Invitrogen, # 00-4958-02). Images were obtained from a Spinning-disk W1 (Yokogawa) or Spinning-disk X1 (Yokogawa) and viewed in Fiji Image J (v.2.16.0).

### Image Analysis

Neurite network analysis was conducted using a custom macro developed for the Fiji software^97^, incorporating functionalities from the Bio-Formats^98^, Analyse Skeleton^99^ and Local Thickness^100^ plugins. Code is openly accessible at: https://github.com/orion-cirb/Axon_Skel_Analyzer.

Nuclei were segmented and counted in the DAPI channel using the following sequence of operations: maximum intensity z-projection, median filtering (radius = 10 pixels), Huang automatic thresholding, hole filling, watershed splitting, object labelling, and filtering based on morphological criteria (minimum area = 40 µm², minimum circularity = 0.7). Cell bodies were segmented in the ORF1p channel by successively applying sum slices z-projection, median filtering (radius = 50 pixels), Huang thresholding, morphological opening (20 iterations) and dilation (5 iterations), hole filling, and size filtering (minimum area = 80 µm2).

The TUJ1 channel was segmented via maximum intensity z-projection, background subtraction (rolling ball radius = 50 pixels), median filtering (radius = 2 pixels), Huang thresholding, morphological closing (4 iterations), an additional median filtering (radius = 2 pixels), selective hole filling (holes < 10 µm²), and size filtering (minimum area = 20 µm2). To isolate neurites, cell bodies segmented from the ORF1p channel were subtracted from the TUJ1 channel binary mask. The resulting neuronal processes were then skeletonized and branches shorter than 4□µm were excluded to minimize artifacts. This processing pipeline enabled local thickness analysis to quantify the distribution of branch diameters.

The analysis of ORF1p intensity after ORF1p-LOF in LUHMES cells was performed using the Nuclei_ORF1p plugin^19^ developed for the Fiji software. This tool enabled quantification of ORF1p signal specifically in the nucleus versus the cytoplasm of cells.

### Subcellular fractionation

Subcellular fractionation was performed with 10^6^ cells using the Subcellular Protein Fractionation kit (Thermo Fisher Scientific, # 78840) according to manufacturer’s recommendations. For cytoplasm and nucleus fractionation, cells were washed in PBS and resuspend in lysis buffer (10mM HEPES ph7.9, 40mM KCL, 5% glycerol, 3mM MgCl2, 0.2% NP40, EDTA-free protease inhibitor cocktail and phosphatase inhibitors) and incubated for 30 min at 4°C. Lysates were passed 20 times in a Dounce homogenizer (loose) and then left for 30 min on a wheel at 4°C. After centrifugation at 4°C at 500g for 5 min, the supernatants were kept as the cytosolic fraction, and the pellets are washed once in lysis buffer and resuspend in nuclear lysis buffer (20mM HEPES ph7.9, 420mM NaCl, 25% glycerol, 1.5mM MgCl2, EDTA-free protease inhibitor cocktail and phosphatase inhibitors) and rotated for 30 minutes on a wheel at 4°C. Finally, lysates were centrifuged for 10 min at 14000g at 4°C and the supernatant was collected as nuclear extract.

### RNA extraction

3.5 x 10^5^ cells were scrapped in TRIzol reagent (Invitrogen, # 15596026) and total RNA was extracted using Direct-zol™ RNA Microprep (Zymo, # R2062) according to manufacturer’s instructions. All samples were subjected to DNAse I treatment during extraction.

### RT-qPCR

300 ng RNA was subject to reverse transcription using All-In-One 5X RT MasterMix kit (Abm, # G592). qPCR was performed in duplicate for each sample and each gene with SsoAdvanced Universal SYBR Green (BioRad, #1725274) with the following protocol: a first heat up at 95°C, followed by the denaturation step at 95°C, annealing at 57°C and finally extension at 72°C for a total of 40 cycles, on a CFX Opus 384 Real-Time PCR System (BioRad, # 12011452). qPCR was analysed by CFX Maestro 2.0 software (v5.0.021.0616) (BioRad) and each gene of interest was normalized to three housekeeping genes: ENO2, TBP and HPRT.

Following primers were used: HPRT (F: CAGCCCTGGCGTCGTGATTAGT; R: CCAGCAGGTCAGCAAAGA AT); TBP (F: CAGCATCACTGTTTCTTGGCGT; R: AGATAGGGATTCCGGGAGTCAT); ENO2 (F: CGTTACTTAGGCAAAGGTGTCC; R: CTCCAGCATCAGGTTGTCCAGT); DLG5 (F: GCCTTCAGAGTCTGCCTTGT; R: CAGGCTGGAAGACAGGAAGG); NEGR1 (F: TTAAGAGGCAGGCTCGCATT; R: TACCAACGTGACACAGGAGC); EFNA5 (F: AGCGGTCCATTTGGAGAGTG; R: ATGAGGACTCCGTCCCAGAA); GABRB3 (F: GCCTTGGCTGTCTTTTCTGC; R: ACCTTATGGGCTGCTTCGTC).

### Immunoprecipitation and sample preparation for mass spectrometry

Magnetic beads were coupled with ORF1p (MN9D: Abcam, #ab245124) (LUHMES: Abcam, #ab245249) or IgG rabbit (Abcam, #ab172730) antibodies using the Dynabeads® Antibody Coupling Kit (Thermo Fisher Scientific, # 14311D) according to the manufacturer’s recommendations. After two washes in DPBS, cells were scraped in lysis buffer (10mM Tris HCl pH:8, 150mM NaCl, 0,5% NP40, protases and phosphatase inhibitors), incubated on ice for 15 min and sonicated (Bioruptor UCD-200) for 15 min (30s on/ 30s off) on ice at medium power. Lysates were incubated with coupled beads and diluted in wash buffer (10 mM Tris HCl pH8, 150 mM NaCl, protases and phosphatase inhibitors) overnight on a wheel at 4°C. Samples were washed 4 times with wash buffer and were resuspended in 100 μl of 25 mM NH4HCO3. IP samples were then washed twice with 100 µL of 25 mM NH4HCO3 (ABC) and proteins were trypsin/lysC (0.2 µg, Promega) digested directly on beads in a total volume of 100µL of 25 mM ABC for one hour at 37°C while vortexed at 1000rpm. Samples were then cleaned on homemade C18 StageTips (AttractSPE Disk Bio C18-100.47.20 Affinisep), before being eluted using 40/60 CH3CN/H2O□+□0.1% formic acid and vacuum concentrated to dryness.

### LC-MS/MS Analysis

LUHMES liquid chromatography was performed with an RSLC nano system (Ultimate 3000, Thermo Scientific) coupled online to an Orbitrap Exploris 480 mass spectrometer (Thermo Scientific). Peptides were trapped on a C18 column (75 μm inner diameter × 2 cm; nanoViper Acclaim PepMap^TM^ 100, Thermo Scientific) with buffer A (2/98 CH3CN/H2O in 0.1% formic acid) and separation was performed on a 50 cm x 75 μm C18 column (nanoViper Acclaim PepMap^TM^ RSLC, 2 μm, 100Å, Thermo Scientific) with a linear gradient of 3% to 29% buffer B (100% CH3CN in 0.1% formic acid, A’: 100% H2O in 0.1% formic acid) at a flow rate of 300 nL/min over 91 min. For Data dependent acquisition (DDA), MS full scans were performed in the ultrahigh-field Orbitrap mass analyser in ranges m/z 375–1500 with a resolution of 120□000 at m/z 200. The top 20 intense ions were subjected to Orbitrap for further fragmentation via high energy collision dissociation (HCD) activation and a resolution of 15□000 with the intensity threshold kept at 1.3 × 10^5^. We selected ions with charge state from 2+ to 6+ for screening. Normalized collision energy (NCE) was set at 27 and the dynamic exclusion of 40s.

MN9D LC-MS/MS analysis was performed, as previously, in data independent acquisition (DIA) mode and by using a linear gradient of 2% to 30% buffer B. MS full scans were performed in the ultrahigh-field Orbitrap mass analyser in range m/z 375– 1500 (120□000 resolution; IT 100 ms; AGC 300%). 40 DIA scan of 15 Da isolation window with 1 Da overlap between 400 to 1000 m/z (15 000 resolution; NCE 25; AGC 3000; IT auto) were set.

For protein identification, LUHMES DDA data were searched against the Homo Sapiens (UP000005640_9606) UniProt database using Sequest HT through proteome discoverer (PD version 2.4). Enzyme specificity was set to trypsin and a maximum of two miss cleavages sites were allowed. Oxidized methionine, Met-loss, Met-loss-Acetyl and N-terminal acetylation were set as variable modifications. Maximum allowed mass deviation was set to 10 ppm for monoisotopic precursor ions and 0.02 Da for MS/MS peaks.

MN9D DIA data were searched against the Mus Musculus (UP000000589) Uniprot database using Spectronaut v18.6 (Biognosys) by directDIA+ analysis using default search settings. Enzyme specificity was set to trypsin and a maximum of two missed cleavage site were allowed. N-terminal acetylation and oxidation of methionine were set as variable modifications.

The resulting files were further processed using myProMS v3.10.0 (https://github.com/bioinfo-pf-curie/myproms)^101^. FDR calculation used Percolator^102^ and was set to 1% at the peptide level for the LUHMES data. The label free quantification was performed by peptide Extracted Ion Chromatograms (XICs) and DDA were reextracted by conditions and computed with MassChroQ version 2.2.21^103^ (LUHMES data). For protein quantification, XICs from proteotypic peptides shared between compared conditions (TopN matching) with missed cleavages were used. Median and scale normalization at peptide level was applied on the total signal to correct the XICs for each biological replicate (N=5). To estimate the significance of the change in protein abundance, a linear model (adjusted on peptides and biological replicates) was performed, and a two-sided t-test was applied on the fold change estimated by the model. The p-values were adjusted using the Benjamini–Hochberg FDR procedure.

The mass spectrometry proteomics raw data have been deposited to the ProteomeXchange Consortium via the PRIDE^104^ partner repository under the accession number PXD066047.

### RNA sequencing and analysis

Four samples for each condition (siRNA-Ctrl and siRNA-Orf1) were sequenced (Novogene Europe). Quantity and quality were assessed using Nanodrop and Bioanalyzer RNA 6000 Pico Kit (Agilent Technologies, # 5067-1513). Libraries including rRNA removal were generated by RNA Library Prep Kit for Illumina following manufacturer’s recommendations and sequenced as pair-end (PE) 150 bp using Novaseq x plus sequencer. Mapping was performed by Hisat2 (v2.0.5) on human genome (hg38/GRCh38 assembly). Differential enrichment analysis was done with DESeq2 (v1.46.0) and data were normalized using DESeq2 scaling and size factors method. Genes where the sum of reads from all samples was under ten reads were removed. Genes with no reads for at least 6 samples on 8 were discarded. Volcano plot was generated by considering as significatively upregulated (264 genes), genes with log2 fold change ≥ 0.5 and *p*□≤ 0.01; significatively downregulated (698 genes), genes with log2 fold change ≤ −0.5, *p* ≤ 0.01). Another more stringent threshold was fixed for defining downregulated genes (log2 fold change ≤ −0.5, *p* ≤ 0.005), and among those, genes forming a network involved in synapse GO (50 genes) were visualized by STRING database (v.12.0). Gene annotations were retrieved from hg38 RefSeq_all and long genes were defined as > 100kB. Kyoto Encyclopedia of Genes and Genomes (KEGG) pathway analysis was conducted with R function enrichKEGG from clusterProfiler (v.4.14.4). Gene Ontology (GO) analysis was executed using the PANTHER GO database (release 19.0). p-values were adjusted using the Benjamini-Hochberg method.

### Human Samples

Cingulate gyrus brain tissue from control (n=5) and Parkinson disease patients (n=5) were provided by the Biobank Neuro-CEB and conserved at −80°C. Tissues underwent one manual physical shock and were then disposed in precooled Tissue-Tube bags (Covaris) and dry cryo-pulverized with a level 6 impact using the CryoPrep Dry Pulveriser (Covaris). All steps were done on dry ice.

### Isolation and sorting of nuclei (FANS)

Nuclei from pulverized tissues of human patients were resuspend in homogenization buffer (HB) (0.25M sucrose, 0.5mM spermidine, 0.15nM spermine, NP40 0.3%, protease inhibitors, 1mM DTT). Tissues were grinded mechanically using a Dounce homogenizer and passed through a 40μm cannula. The recovered cells were then put on an OptiPrep gradient (Sigma, # D1556) and centrifuged for 15 minutes at 10000g. Pellets were dried and put in PBS spermine 0.15nM. Human nuclei were then labelled 1h on a wheel at 4°C using the following antibodies: NeuN PE (Merck, # FCMAB317PE), ORF1p AF647 (Abcam, # ab314927) and control antibodies: IgG1 mouse-PE (Beckman Coulter, # A07796) and IgG rabbit-AF647 (Abcam, # ab199093). Nuclei sorting was performing using a FACSARIA II cell sorter (Beckton Dickinson) equipped with three lasers (405, 488 and 640nm), a 70µm nozzle and FACSDiva software (version 6.1.2). Nuclei were detected according to their forward scattering light (FSC) and gated using forward and side scatter light signals. Aggregates were excluded by forward scatter area versus height pulse measurement. Finally, PE and AF647 fluorescence were respectively measured through a 585/42 and 660/20nm bandpass filters. P3 corresponds to all nuclei (defined as ORF1p-high) which fluoresce beyond the maximum intensity of nuclei in the control (non-relevant antibody that fluoresces in the same channel as ORF1 (x-axis, 647nm)), while P4 is everything below this threshold (ORF1p-low nuclei). As in the control (non-relevant antibody that fluoresces in the same channel as NeuN (y-axis, NeuN-PE)), autofluorescent nuclei form a diagonal, we excluded nuclei in this range and defined as NeuN-positive all nuclei above this autofluorescent window. 50 000 nuclei were recorded. Nuclei were collected in PBS buffer in Eppendorf tubes.

### ATAC sequencing and analysis

After FANS, 50,000 nuclei per condition were centrifuged at 4°C, 500g for 10 min. ATAC-seq was then performed using a commercial kit (Diagenode Cat. No.C01080006 for human tissue and Cat. No.C01080001 for LUHMES) according to the instructions of the manufacturer. Profiles of libraries were verified using the Bioanalyzer High Sensitivity DNA Kit (Agilent Technologies). The library pool was quantified by qPCR using the KAPA library quantification kit (Roche). Sequencing was carried out on the NovaSeq 6000 instrument from Illumina using paired-end 2 x 100 bp sequencing. ATAC-seq data were analyzed using the Curie Institute ATAC-seq pipeline (v1.02, DOI: 10.5281/zenodo.7576558 and available at https://github.com/bioinfo-pf-curie/ATAC-seq/releases/tag/v1.0.2). Briefly, sequencing reads were aligned after adapter removal on the Human reference genome (hg38) with bwa-mem. Aligned reads were filtered to remove mitochondrial reads, PCR duplicates, singleton and reads aligned with a mapping quality lower than 20. Genomic sites from the ENCODE blacklist regions were also discarded from the analysis. The peak calling was then performed using the MAC2 software with the following parameters “-f BAMPE --nomodel -q 0.01”. DeepTools (v3.5.4) command multiBamSummary was used to count reads from all peaks. All peaks were kept and peaks with the sum of reads from all samples smaller than ten reads were removed. Differential accessible peaks were determined with DESeq2 (v1.46.0). For LUHMES, we compared siRNA-Ctrl (n=2) and siRNA-ORF1 (n=2) and peaks were considered as significant with a p-value□≤ 0.05 and log2FC□≥ and ≤ 0.5. For human samples (NeuN-= 4 samples, NeuN+ ORF1p low = 6 samples and NeuN+ ORF1p high = 6 samples), peaks were considered as significant with a p-value□≤ 0.01 and log2FC□≥ and ≤ 1. Peaks were annotated to the nearest gene with the function annotatePeak from ChIPseeker R package (v.1.42.1), using *TxDb.Hsapiens.UCSC.hg38.knownGene* and TSSregion set to ±□3□kb. Motif enrichment analysis was performed with HOMER (v5.1), using findMotifsGenome.pl, with default parameters. GO analysis was conducted on the PANTHER GO database (release 19.0). Significative DAR association with known chromatin states was investigated using available ChromHMM annotations (retrieved from Roadmap= https://pmc.ncbi.nlm.nih.gov/articles/PMC8734071/) from adult cingulate gyrus (ATAC human brain) and substantia nigra (ATAC LUHMES). 71 significant genes were overlapping between ATAC-seq in human post mortem tissues and LUHMES. GO BP analysis of those proteins were carried out in R using the clusterProfiler (v.4.14.4). package and represented as a GOChord plot (package GOplot v.1.0.2). Bigwig tracks were visualized using the Integrated Genome Viewer (IGV) browser (v2.19.1).

### Statistics

Statistical analyses were done with Graph Pad PRISM software (v10.4.1). The significance threshold was defined as p<0.05 except stated otherwise. We assessed the distribution of differentially accessible regions (DARs) across genomic features or regulatory regions using two-sided Fisher’s exact tests. To ensure statistical robustness, categories with low total counts (<5 DARs) were excluded from these analyses. False discovery rate (FDR) correction with reported q-values was applied to p-values of the remaining comparisons using the Benjamini–Hochberg procedure (Q =1%). Gene biotype and long gene overrepresentation analyses were done using a Chi-Square test with observed = number of feature counts in ORF1p-LOF and expected = all RNA-seq transcript counts.

## Acknowledgements

SS acknowledges Université PSL for a Ph.D. fellowship and was enrolled with the Ecole Doctorale ED3C. This work was supported by the Agence Nationale de la Recherche (National French Agency for Research; ANR-20-CE16-0022 NEURAGE), the “Fondation du Collège de France” (to J.F.), the “Fondation NRJ/Institut de France” (to J.F.), the “Fondation Alzheimer” (to J.F.) and the “Fédération pour la Recherche sur le Cerveau” (FRC, to J.F.). High-throughput sequencing was performed by the ICGex NGS platform of the Institut Curie supported by the grants ANR-10-EQPX-03 (Equipex) and ANR-10-INBS-09-08 (France Génomique Consortium) from the “Agence Nationale de la Recherche” (“Investissements d’Avenir” program), by the ITMO-Cancer Aviesan (Plan Cancer III) and by the SiRIC-Curie program (SiRIC Grant INCa-DGOS-465 and INCa-DGOS-Inserm_12554). Data management, quality control and primary analysis were performed by the Bioinformatics platform of the Institut Curie. The LSMP thanks Patrick Poullet from the bioinformatics platform of the Institut Curie U1331 for the continuous development of myProMS. We thank the “The Brainbank Neuro-CEB Neuropathology Network” for the human *post-mortem* tissues. The Neuro-CEB Neuropathology network includes: Dr Franck Letournel (CHU Angers), Dr Marie-Laure Martin-Négrier (CHU Bordeaux), Dr Maxime Faisant (CHU Caen), Pr Catherine Godfraind (CHU Clermont-Ferrand), Pr Claude-Alain Maurage (CHU Lille), Dr Vincent Deramecourt (CHU Lille), Dr Mathilde Duchesne (CHU Limoges), Dr David Meyronnet (CHU Lyon), Dr André Maues de Paula (CHU Marseille), Pr Valérie Rigau (CHU Montpellier), Dr Fanny Vandenbos-Burel (Nice), Pr Charles Duyckaerts (CHU PS Paris), Pr Danielle Seilhean (CHU PS, Paris), Dr Susana Boluda (CHU PS, Paris), Dr Isabelle Plu (CHU PS, Paris), Dr Serge Milin (CHU Poitiers), Dr Dan Christian Chiforeanu (CHU Rennes), Dr Florent Marguet (CHU Rouen), Dr Béatrice Lannes (CHU Strasbourg). We would also like to thank the following patient organisations which support the Neuro-CEB brainbank: ARSLA, CSC, France DFT, Fondation ARSEP, Fondation Vaincre Alzheimer and France Parkinson. We also thank the Fondation Bettencourt Schueller for their support. We especially thank Ariel Di Nardo and Alain Prochiantz for their continuous support.

## Contributions

S.S. carried out the majority of the experimental work and collected and analyzed the data. O.M.B. designed and performed the FANS analysis with M.F. and S.S., carried out the subcellular fractionations with S.S. and prepared the IP samples for mass spectrometry (LUHMES). B.L. carried out the LC-MS/MS experimental work and D.L. supervised MS and MS data analysis which was followed up by S.S and T.B.. H. M. and T.C. set up the script for image analysis, N.S. supervised the bioinformatical analyses done by S.S.. S.S., R.L.J. and J.F interpreted the data, S.S. and J.F. wrote the manuscript with input from all authors. J.F. and R.L.J. conceived and supervised the project.

## Competing interests

The authors declare no competing interests.

## Supplementary Figures

**Suppl. Figure 1:**
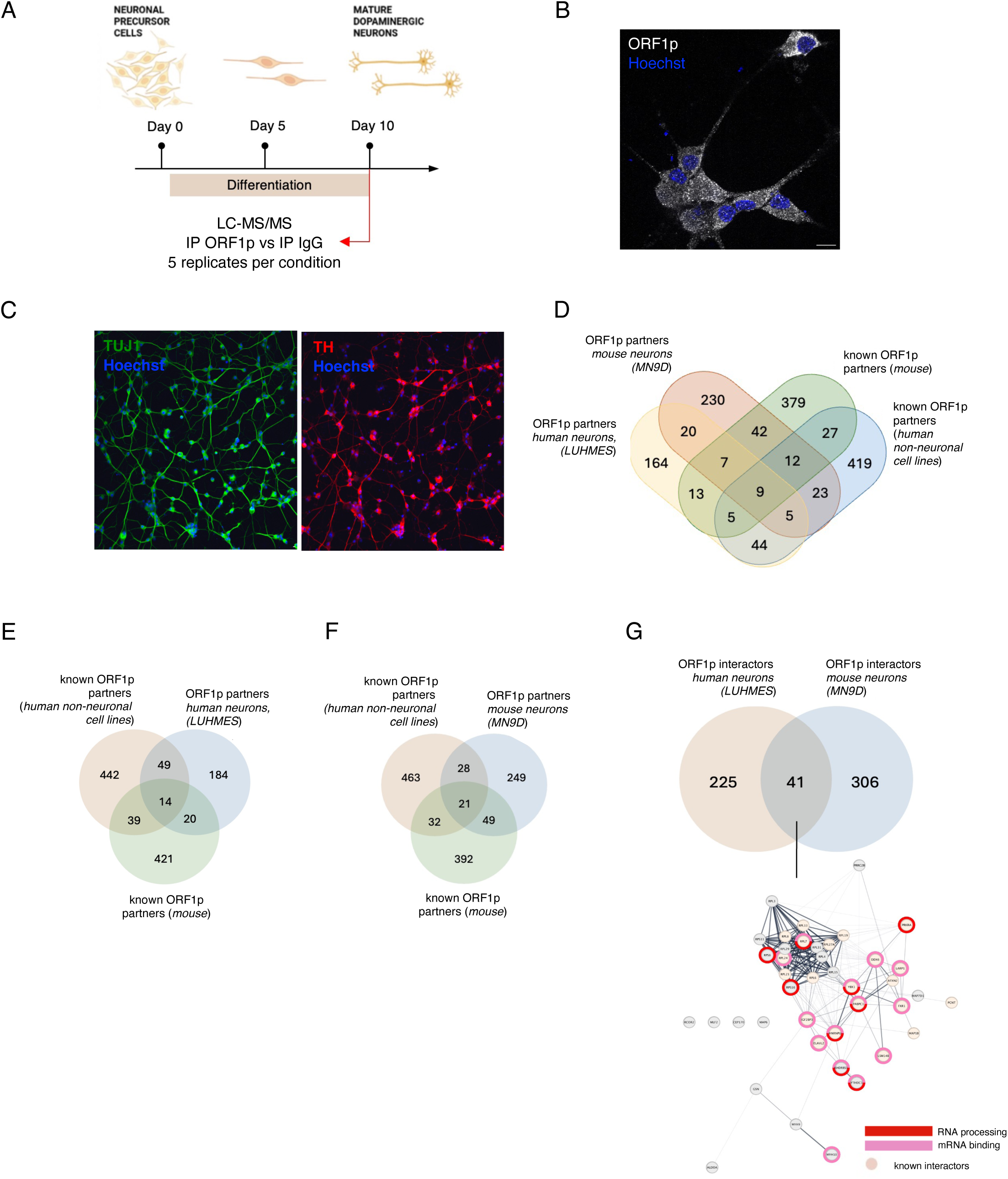
**A.** Scheme of the differentiation protocol of MN9D cells into mature mouse dopaminergic neurons. All experiments (including quantitative mass spectrometry) were performed at day 10 of differentiation. **B.** Immunofluorescence staining of endogenous ORF1p (ab216324) in differentiated MN9D neurons. **C.** Immunofluorescence staining of MN9D at 10 days of differentiation expressing the neuronal marker TUJ1 (green, left) and the dopaminergic marker tyrosine hydroxylase (TH, red, right). **D.** Venn Diagram of proteins identified by quantitative mass spectrometry of immunoprecipitated endogenous ORF1p from mouse dopaminergic neurons (MN9D) and human dopaminergic neurons (LUHMES) compared to mouse and human published ORF1p interactors^17,26–34,63^. **E.** Venn Diagram of proteins identified by quantitative mass spectrometry of immunoprecipitated endogenous ORF1p from human dopaminergic neurons (LUHMES) compared to previously published mouse and human ORF1p interactors^26–34^. **F.** Venn Diagram of proteins identified by quantitative mass spectrometry of immunoprecipitated endogenous ORF1p from mouse dopaminergic neurons (MN9D) compared to previously published mouse and human ORF1p interactors^17,63^. **G.** Common partners of ORF1p in LUHMES and MN9D neurons and their physical interactions identified by quantitative mass spectrometry.

**Suppl. Figure 2:**
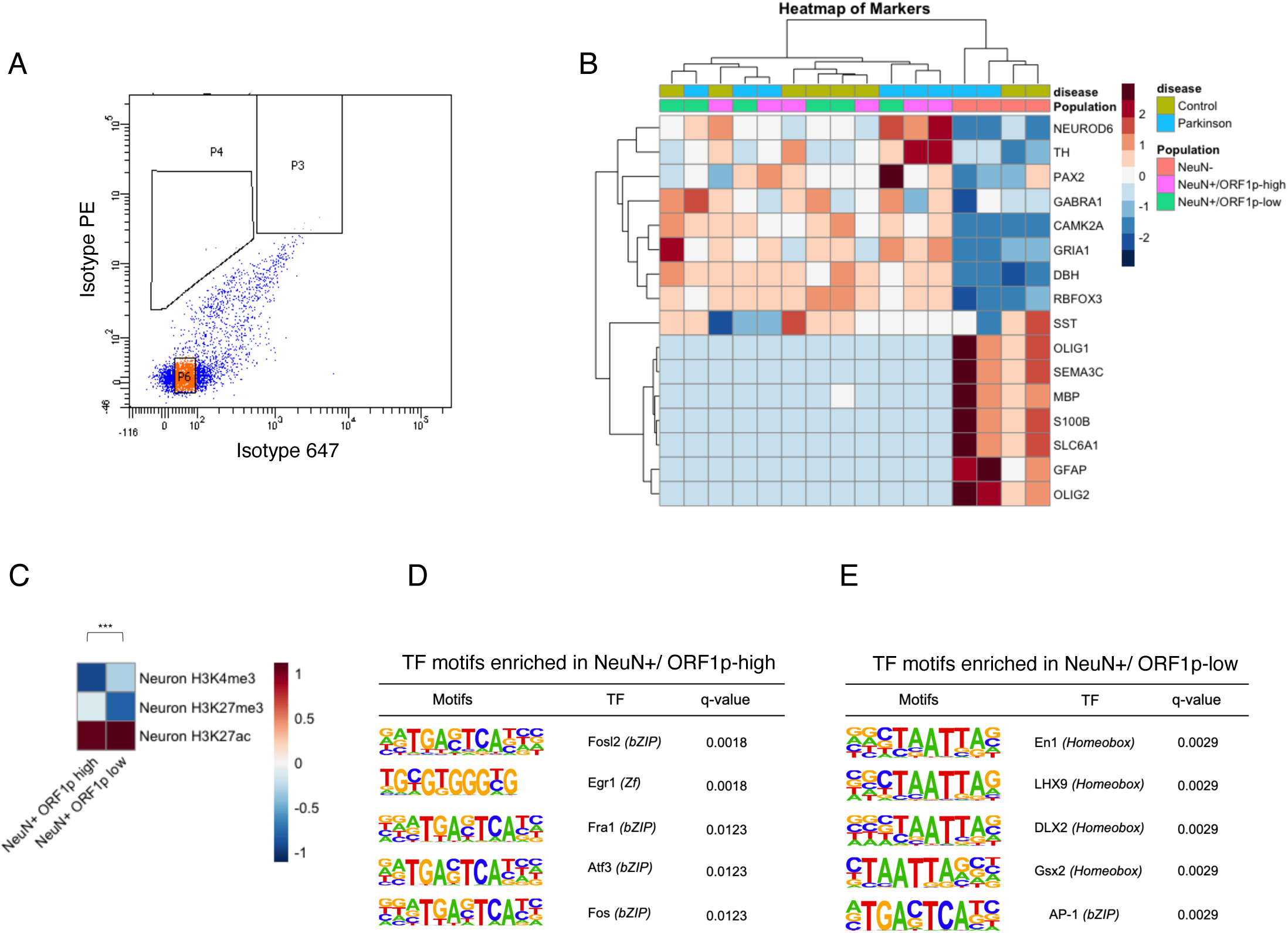
**A.** Representative fluorescent-activated nuclei sorting (FANS) profile showing the isotype controls for the NeuN and the ORF1p antibody used in Figure 3B. **B.** Heatmap of brain cell type markers and ATAC seq promoter peak numbers (<1kB) in NeuN-, NeuN+ ORF1p-low and NeuN+ ORF1p-high populations. NeuroD6, TH, PAX2, GABRA1, CAMK2A, GRIA1, DBH and RBFOX3 (= NeuN) and SST are markers of specific neuronal populations (https://biomarkerres.biomedcentral.com/articles/10.1186/s40364-023-00523-3) and show a high accessibility on promoters in both, ORF1p-high and ORF1p-low nuclei. OLIG1, SEMA3C, MBP, S100B, SLC6A1, GFAP and OLIG2 are glia-cell markers and show higher chromatin accessibility in promoter regions in non-neuronal (NeuN-) nuclei. **C.** Heatmap of differentially accessible regions in ORF1p-high and ORF1p-low nuclei and their overlap with the histone marks H3K4me3, H3K27me3 and H3K27ac. Histone post-translational modification profiles were retrieved from a previously published dataset^105^ **D-E**. Motif enrichment analysis of significant DARs in ORF1p-high and ORF1p-low neuronal populations.

**Suppl Figure 3:**
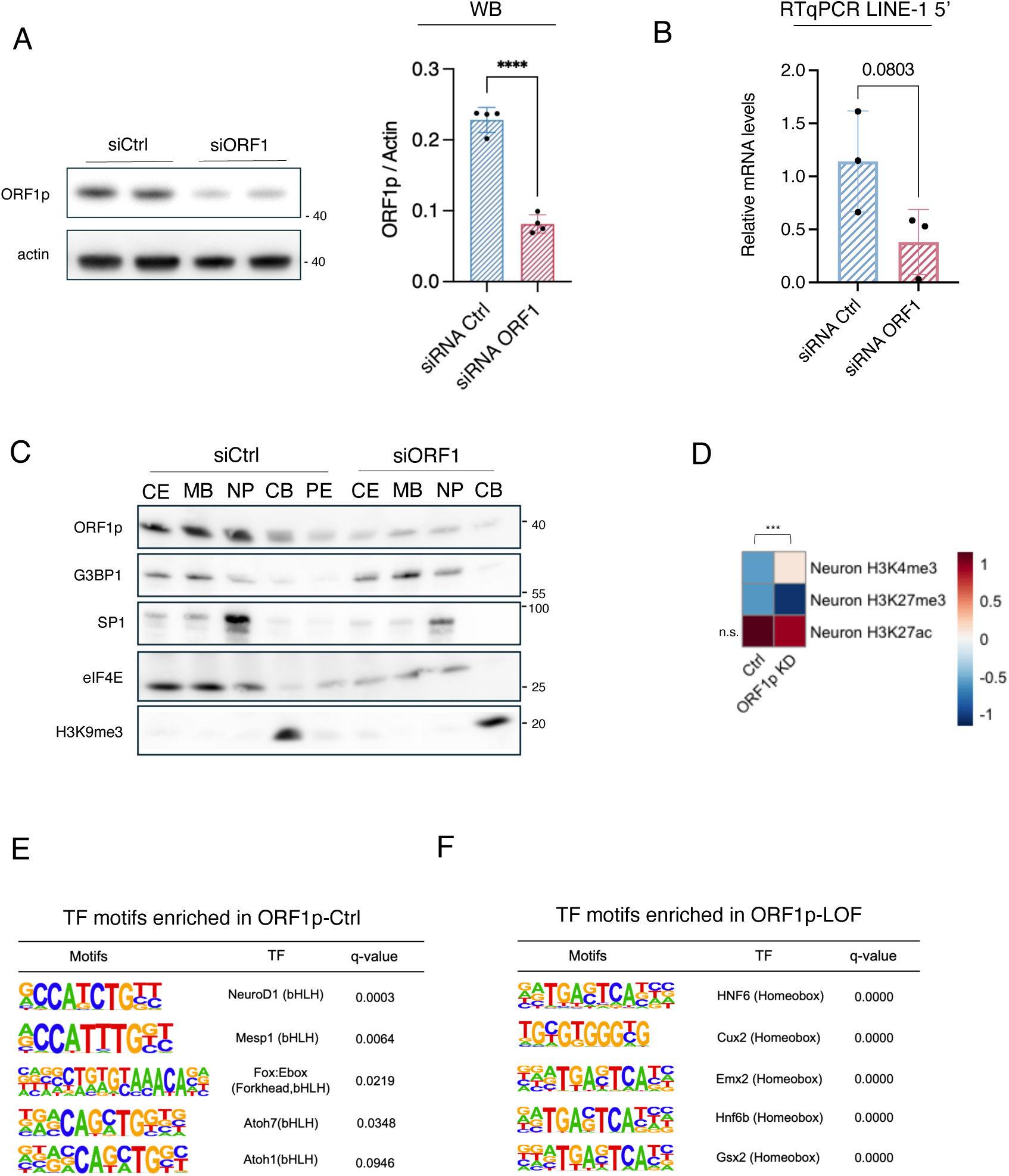
**A.** Western Blot and quantification showing ORF1p-LOF in differentiated LUHMES neurons (n = 4 wells; three independent experiments, day 7 post-differentiation). Two-tailed Mann–Whitney *U test*. Data are represented as mean□±□SEM. **B.** Subcellular fractionation of control (siRNA-Ctrl) or ORF1p-LOF (siRNA ORF1) LUHMES neurons. Shown are the cytoplasm (CE; marker: eIF4E), the membrane-bound fraction (MB), the nucleoplasm (NP; marker: SP1), the chromatin-bound fraction (CB; marker: H3K9me3) and the pellet (PE). **C.** LINE-1 5’UTR RT-qPCR of control (siRNA-Ctrl) and ORF1p-LOF (siRNA ORF1) LUHMES neurons. **D.** Heatmap of differentially accessible regions in Ctrl and ORF1p-LOF nuclei and their overlap with the histone marks H3K4me3, H3K27me3 and H3K27ac. Histone post-translational modification profiles were retrieved from a previously published dataset^105^. **E-F**. Motif analysis of significant DARs identified in ATAC-seq in Control and ORF1p-LOF human dopaminergic neurons (LUHMES).

**Suppl Figure 4:**
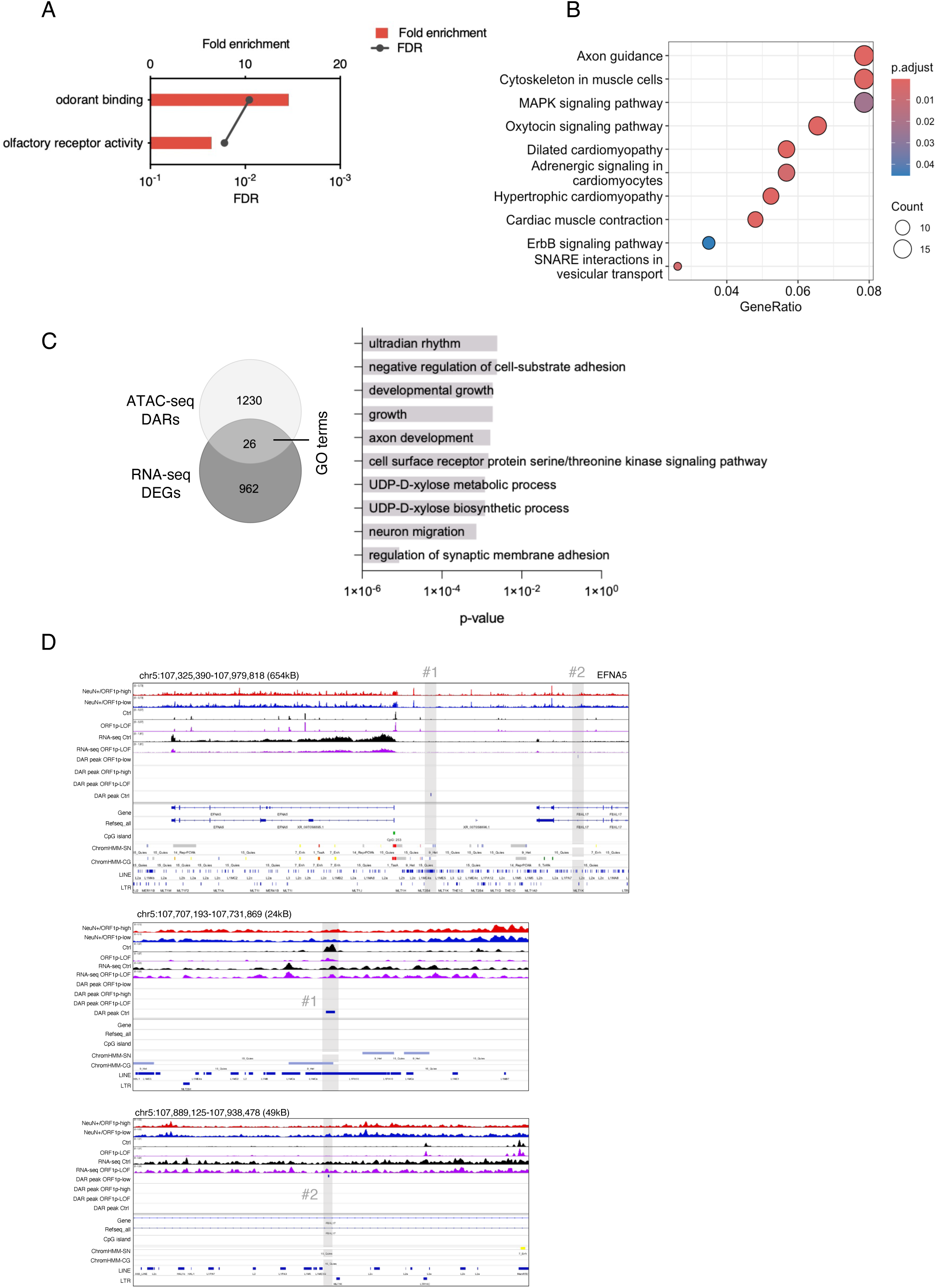
**A.** GO analysis of significantly upregulated transcripts in ORF1-LOF vs Ctrl neurons. FDR was calculated from p-values (Fisher’s exact test) by the Benjamini-Hochberg procedure. MF: Molecular Function. **B.** Kyoto Encyclopedia of Genes and Genomes (KEGG) pathway analysis of significantly downregulated genes upon ORF1p-LOF. P-value was calculated using one-sided Fisher’s exact test and the p-value was adjusted for multiple testing by the Benjamini-Hochberg method. **C.** Venn diagram of common differentially accessible regions (DARs) in ATAC-seq (1230 total) and differentially regulated genes (DEGs) in RNA-seq (962 total) from human neurons (LUHMES). GO (biological process, BP) of the 26 common genes. **D.** IGV track of an example of the neuron-specific gene EFNA5 with DARs in human post-mortem cingulate gyrus (ORF1p-high versus ORF1p-low) and human LUHMES neurons (Ctrl versus ORF1p-LOF). EFNA5 was significantly downregulated upon ORF1p-LOF.

**Suppl. Table 1.** ORF1p protein partners in LUHMES and MN9D as identified by quantitative mass spectrometry.

**Suppl Table 2.** Metadata of post-mortem cingulate gyrus tissues used in FANS/ATAC-seq.

| Diagnosis | Gender | Age | PMI (hrs) | Braak Stage |
| --- | --- | --- | --- | --- |
| PD | F | 62 | - | - |
| PD | F | 79 | - | II |
| PD | M | 83 | 20 | II |
| non-PD | F | 92 | 21 | I |
| non-PD | M | 84 | 15 | - |
| non-PD | M | 73 | 10 | II |

PD: Parkinson disease
PMI: post-mortem interval

**Suppl Table 3.** Characteristics of significantly enriched “differentially accessible regions” more accessible (DAR+) in neurons with either high or low nuclear ORF1p content identified by ATAC-seq in human post-mortem cingulate gyrus neurons and human dopaminergic neurons in culture (LUHMES). In red overlapping characteristics in both experimental systems are highlighted.

| human brain |  | ORF1p-high | ORF1p-low |
| --- | --- | --- | --- |
|  | genomic region | distal intergenic | intronic |
|  | functional feature | heterochromatin | flanking active TSS |
|  |  | weak repressed polycomb | enhancer |
|  | histone profile | H3K27me3 | H3K4me3, H3K27ac |
|  | TF binding sites | bZIP, ZF | Homeobox, bZIP |

differentially **more** accessible regions (DAR+)
| LUHMES |  | Control | ORF1p-LOF |
| --- | --- | --- | --- |
|  | genomic region |  |  |
|  | functional feature | repressed polycomb | flanking active TSS |
|  |  | weak repressed polycomb | enhancer |
|  |  | ZNF/Repeats |  |
|  | histone profile | H3K27me3 | H3K4me3 |
|  | TF binding sites | bHLH | Homeobox |

in red= overlapping terms in both ATAC-seq experiments

## Notes

### Competing Interest Statement

The authors have declared no competing interest.

